# Novel enzyme for dimethyl sulfide-releasing in bacteria reveals a missing route in the marine sulfur cycle

**DOI:** 10.1101/2020.10.29.360743

**Authors:** Chun-Yang Li, Xiu-Juan Wang, Xiu-Lan Chen, Qi Sheng, Shan Zhang, Peng Wang, Mussa Quareshy, Branko Rihtman, Xuan Shao, Chao Gao, Fuchuan Li, Shengying Li, Yin Chen, Yu-Zhong Zhang

## Abstract

Dimethylsulfoniopropionate (DMSP) is an abundant and ubiquitous organosulfur molecule and plays important roles in the global sulfur cycle. Cleavage of DMSP produces volatile dimethyl sulfide (DMS), which has impacts on the global climate. Multiple pathways for DMSP catabolism have been identified. Here we identified yet another novel pathway, the ATP DMSP lysis pathway. The key enzyme, AcoD, is an ATP-dependent DMSP lyase. AcoD belongs to the acyl-CoA synthetase superfamily, which is totally different from other DMSP lyases, showing a new evolution route. AcoD catalyses the conversion of DMSP to DMS by a two-step reaction: the ligation of DMSP with CoA to form the intermediate DMSP-CoA, which is then cleaved to DMS and acryloyl-CoA. The novel catalytic mechanism was elucidated by structural and biochemical analyses. AcoD is widely distributed in many bacterial lineages including Alphaproteobacteria, Betaproteobacteria, Gammaproteobacteria and Firmicutes, revealing this new pathway plays important roles in global DMSP/DMS cycles.

## Main

The organosulfur molecule dimethylsulfoniopropionate (DMSP) is produced in massive amounts by many marine phytoplankton, macroalgae, angiosperms, bacteria and animals (***Curson et al., 2018; Stefels, 2000; Otte et al., 2004; Curson et al., 2017; Raina et al., 2013***), and plays important roles in the global sulfur cycle (***Kiene et al., 2000; Charlson et al., 1987***). DMSP can function as an antioxidant, predator deterrent, cryoprotectant or chemoattractant, and may enhance the production of quorum sensing molecules (***Sunda et al., 2002; Wolfe et al., 1997; Karsten et al., 1996; Seymour et al., 2010; Johnson et al., 2016***). Environmental DMSP can be taken up as a carbon and/or sulfur source by diverse bacteria (***Curson et al., 2011***). Bacteria can also degrade DMSP and release volatile dimethyl sulfide (DMS) and methanethiol (MeSH) (***Reisch et al., 2011a***). DMS is the primary natural source of sulfur transferred from oceans to the atmosphere (***Andreae, 1990***), which may participate in the formation of cloud condensation nuclei, and influence the global climate (***Vallina et al., 2007***).

It is known that bacteria can catabolize DMSP via two different pathways, the demethylation pathway and the lysis pathway (***Curson et al., 2011***). Recently, an oxidation pathway for DMSP catabolism was reported (***Thume et al., 2018***). The nomenclature of these pathways is based on the reaction type of the enzyme involved in the first step of DMSP catabolism. In the demethylation pathway, DMSP demethylase DmdA first demethylates DMSP to produce methylmercaptopropionate (MMPA) (***Howard et al., 2006***), which is further catabolized by MMPA-CoA ligase (DmdB), MMPA-CoA dehydrogenase (DmdC) and methylthioacrylyl-CoA hydratase (DmdD/AcuH) into MeSH and acetaldehyde (Fig. 1) (***Reisch et al., 2011b; Bullock et al., 2017; Shao et al., 2019***). In the lysis pathway, eight different lyases (DddD, DddP, DddL, DddQ, DddW, DddK, DddY and Alma1) were identified, cleaving DMSP to produce DMS and acrylate/3-hydroxypropionate (3-HP), which are further metabolized by other enzymes (***Curson et al., 2011; Johnston et al., 2016***). Among these DMSP lyases, only DddD catalyzes an acetyl-CoA-dependent CoA transfer reaction, while the others catalyze a direct cleavage of DMSP (***Bullock et al., 2017; Todd et al., 2007; Alcolombri et al., 2014; Lei et al., 2018***). Thus, the pathway initiated by DddD was later termed as the "CoA DMSP lysis pathway" (***Johnston et al., 2016***), which is separated from the "DMSP lysis pathway" initiated by other DMSP lyases (***Johnston et al., 2016***) (Fig. 1). In the oxidation pathway, DMSP is oxidized to dimethylsulfoxonium propionate (DMSOP), which is further metabolized to dimethylsulfoxide (DMSO) and acrylate; however, enzymes involved in this pathway are still unknown (***Thume et al., 2018***) (Fig. 1). Recently, several bacterial isolates were reported to produce DMS from DMSP without known DMSP lyases in their genomes (***Liu et al., 2018; Zhang et al., 2019***), suggesting the presence of novel pathway(s) for DMSP degradation in nature.

**Fig. 1.**
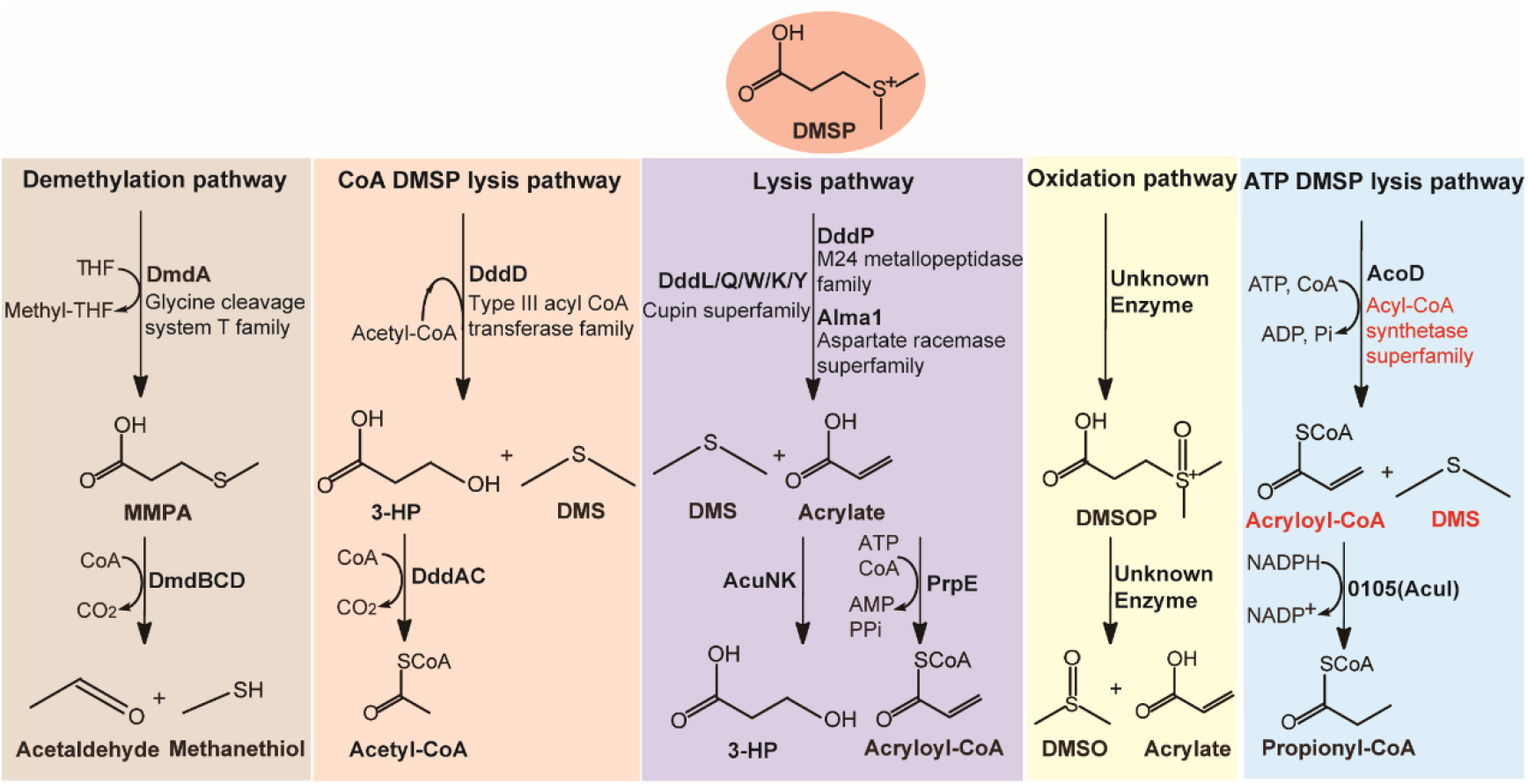
Metabolic pathways for DMSP degradation. Different pathways are shown in different colors. The demethylation of DMSP by DmdA produces MMPA. In the CoA DMSP lysis pathway, the DddD enzyme converts DMSP to 3-HP and DMS, using acetyl-CoA as a CoA donor. In the lysis pathway, DMSP lyase DddP, DddL, DddQ, DddW, DddK, DddY or Alma1 converts DMSP to acrylate and DMS. The oxidation of DMSP produces DMSOP. In our proposed ATP DMSP lysis pathway of this study, DMSP is converted to acryloyl-CoA and DMS via AcoD, with ATP and CoA as cofactors. The protein families of enzymes involved in the first step of each pathway are indicated. The protein family of AcoD and the products of its catalysis are highlighted in red color.

A common feature of previously characterized DMSP metabolic pathways is that the metabolites (*i.e.* MMPA, acrylate, 3-HP) need to be ligated with CoA for further catabolism (Fig. 1) (***Curson et al., 2011; Reisch et al., 2011b***). Thus, it is tempting to speculate that there may be a novel DMSP metabolic pathway, in which DMSP is ligated with CoA first before further catabolism by other enzymes. In this study, we screened DMSP-catabolizing bacteria from Antarctic samples, and obtained a strain *Psychrobacter* sp. D2 that grew on DMSP while producing DMS. Genetic and biochemical results demonstrated that *Psychrobacter* sp. D2 possesses a novel "ATP DMSP lysis pathway" for DMSP catabolism (Fig. 1). The key enzyme in this pathway, AcoD, is an ATP-dependent DMSP lyase which catalyzes a two-step reaction: the ligation of DMSP and CoA, and the cleavage of DMSP-CoA to produce DMS and acryloyl-CoA. We further solved the crystal structure of AcoD and elucidated the molecular mechanism for its catalysis based on structural and biochemical analyses. AcoD is widely distributed in bacteria, being found in both Gram-negative bacteria and Gram-positive bacteria. Our results provide novel insights into the microbial metabolism of DMSP.

## Results

### An unconventional gene cluster involved in DMSP catabolism

With DMSP (5 mM) as the sole carbon source, DMSP-catabolizing bacteria were isolated from five Antarctic samples including alga, sediments and seawaters (Supplementary Fig. 1a, Supplementary Table 1). In total, 175 bacterial strains were obtained (Supplementary Fig. 1b). Among these bacterial strains, *Psychrobacter* sp. D2 grew well in the medium containing DMSP as the sole carbon source (Fig. 2a). Moreover, gas chromatography (GC) analysis showed that *Psychrobacter* sp. D2 could catabolize DMSP and produce DMS (Fig. 2b).

**Fig. 2.**
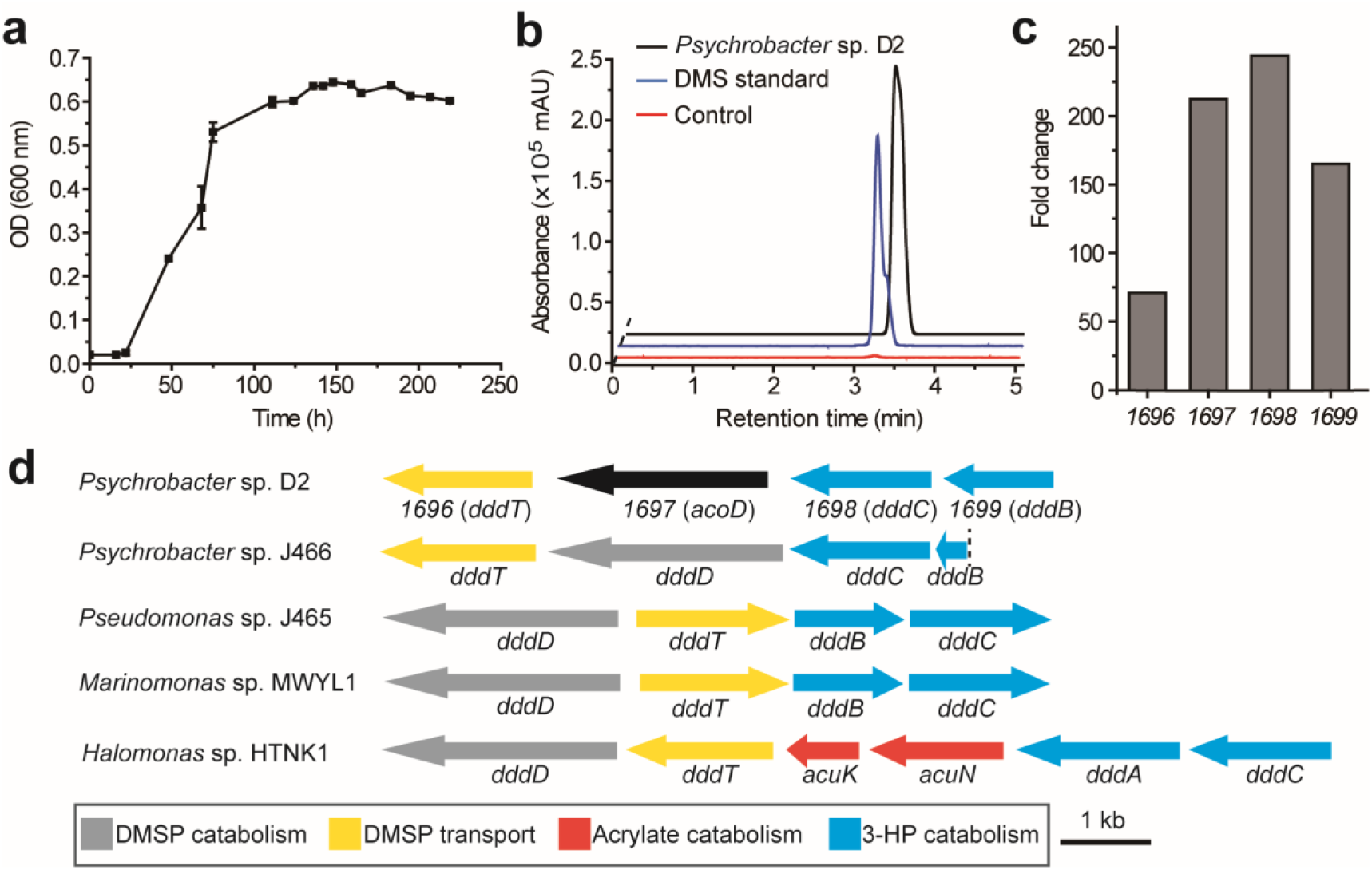
The utilization of DMSP by *Psychrobacter* sp. D2 and the putative DMSP-catabolizing gene cluster in its genome. **a**, The growth curve of *Psychrobacter* sp. D2 on DMSP at 15°C. The error bar represents standard deviation of triplicate experiments. **b**, GC detection of DMS production from DMSP by strain D2. The culture medium without strain D2 was used as the control. The DMS standard was used as a positive control. **c**, Transcriptomic analysis of the putative genes involved in DMSP metabolism in strain D2. The transcriptions of four genes in a cluster were significantly up-regulated during the growth of strain D2 on DMSP. The fold changes were calculated by comparing to the control (transcriptions of these genes during the strain growth on sodium pyruvate). The locus tags of *1696*, *1697*, *1698* and *1699* are H0262_08195, H0262_08200, H0262_08205 and H0262_08210, respectively. **d**, Genetic organization of the putative DMSP-catabolizing gene cluster. Genes from *Psychrobacter* sp. J466, *Pseudomonas* sp. J465, *Marinomonas* sp. MWYL1 and *Halomonas* sp. HTNK1 which were reported to be involved in DMSP catabolism (***Todd et al., 2007; Todd et al., 2010; Curson et al., 2010***) or transport (***Curson et al., 2011***) are shown. The dashed vertical line indicates a breakpoint in *dddB* in the cosmid library of *Pseudomonas* sp. J466 (***Curson et al., 2010***).

To identify the genes involved in DMSP degradation in *Psychrobacter* sp. D2, we sequenced its genome and searched homologs of known DMSP lyases. However, no homologs of DMSP lyases with sequence identity higher than 30% was found in its genome (Supplementary Table 2), implying that this strain may possess a novel enzyme or a novel pathway for DMSP catabolism. We then sequenced the transcriptomes of this strain with/without DMSP as the sole carbon source. Transcriptional data analyses showed that the transcriptions of 4 genes (*1696*, *1697*, *1698* and *1699*) that compose a gene cluster were all significantly upregulated (Fig. 2c) when DMSP was the sole carbon source, which was further confirmed by RT-qPCR analysis (Supplementary Fig. 2). These results suggest that this gene cluster likely participates in the DMSP catabolism in *Psychrobacter* sp. D2.

In the gene cluster, 1696 is annotated as a betaine-carnitine-choline transporter (BCCT), sharing 32% amino acid identity with DddT, the predicted DMSP transporter in *Marinomonas* sp. MWYL1 (***Todd et al., 2007***); 1697 is annotated as an acetate-CoA ligase, and shares 26% sequence identity with the acetyl-CoA synthetase in *Giardia lamblia* (***Sánchez et al., 2000***); 1698 is annotated as an aldehyde dehydrogenase, sharing 72% sequence identity with DddC in *Marinomonas* sp. MWYL1 (***Todd et al., 2007***); and 1699 is annotated as an alcohol dehydrogenase, sharing 65% sequence identity with DddB in *Marinomonas* sp. MWYL1 (***Todd et al., 2007***). DddC and DddB have been reported to be involved in DMSP catabolism (***Todd et al., 2007; Todd et al., 2010***). The pattern of the identified gene cluster *1696*-*1699* in *Psychrobacter* sp. D2 is similar to the patterns of those DMSP-catabolizing clusters reported in *Pseudomonas*, *Marinomonas* and *Halomonas*, in which *dddT, dddB* and *dddC* are clustered with the DMSP lyase gene *dddD* (***Todd et al., 2007; Todd et al., 2010; Curson et al., 2010***) (Fig. 2d). These data strongly suggest that the gene cluster *1696*-*1699* participates in DMSP utilization in *Psychrobacter* sp. D2. However, the sequence identity between 1697 and DddD is less than 15%, suggesting that 1697 is unlikely a DddD homolog. Because the genes adjacent to *1697* in the cluster are deduced to participate in the transport or metabolism of DMSP, we suspected that 1697 is a key enzyme in DMSP degradation in *Psychrobacter* sp. D2, which we termed as AcoD hereafter.

### The essential role of AcoD in DMSP degradation in *Psychrobacter* sp. D2

To identify the possible function of *acoD* in DMSP catabolism, we knocked out this gene from the genome of *Psychrobacter* sp. D2 and constructed the mutant Δ*acoD* (Supplementary Fig. 3). The Δ*acoD* mutant could not grow with DMSP as the sole carbon source, and the complementation of *acoD* via pBBR1MCS-*acoD* to the Δ*acoD* mutant resumed the phenotype of DMSP utilization (Fig. 3a), indicating that *acoD* is essential for strain D2 to utilize DMSP. To further confirm the role of *acoD* in DMSP catabolism, the Δ*acoD* mutant and its complemented strain (Δ*acoD*/pBBR1MCS-*acoD*) were cultured in the marine broth 2216 medium, and then induced by DMSP. The Δ*acoD* mutant produced little DMS, whereas its complemented strain produced substantial amounts of DMS from DMSP (Fig. 3b), which indicated that *acoD* encodes a functional enzyme to degrade DMSP to DMS.

**Fig. 3.**
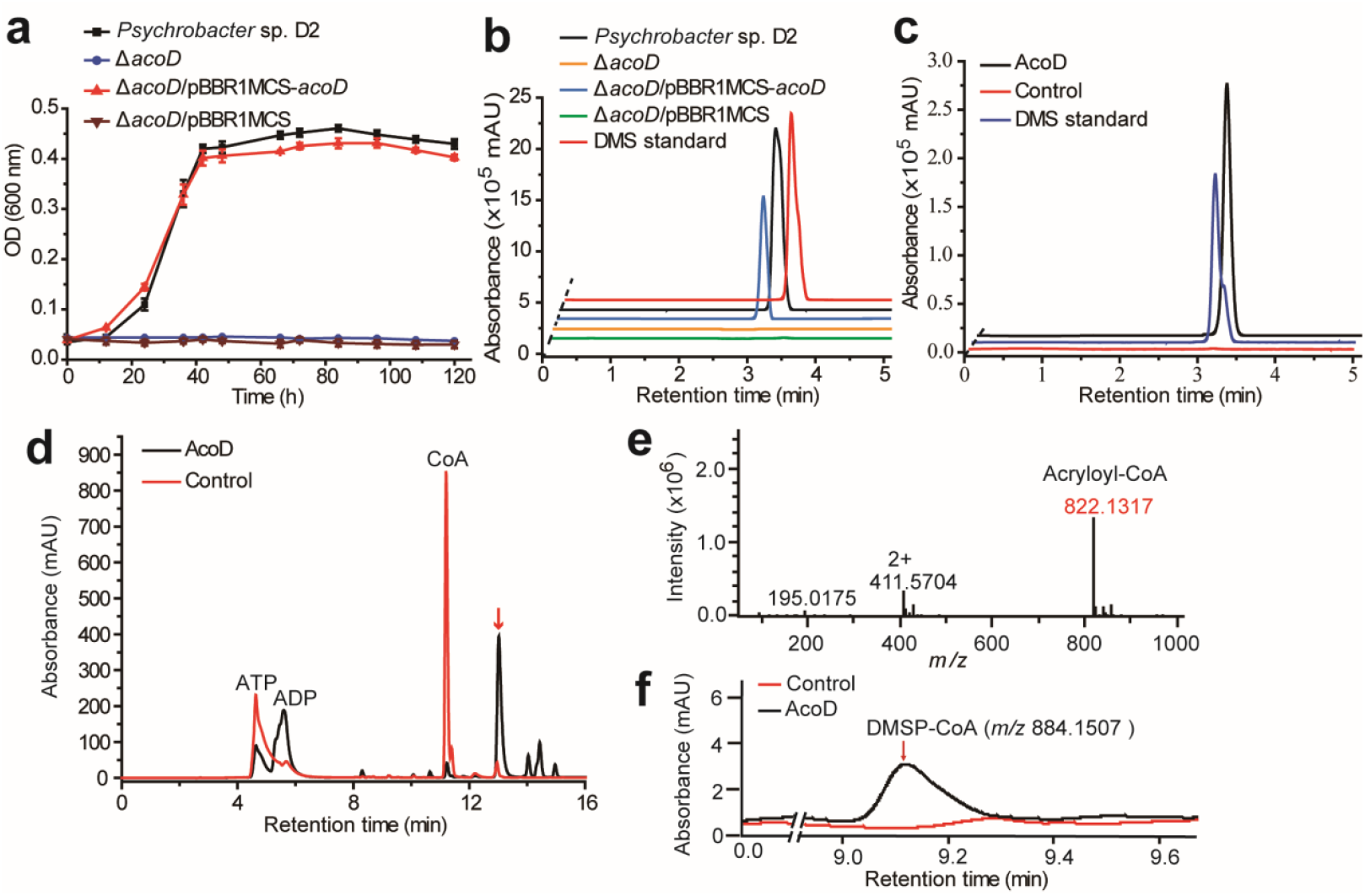
The function of *acoD* of *Psychrobacter* sp. D2 in DMSP metabolism. **a**, Growth curves of the wild-type strain D2, the mutant strain Δ*acoD*, the complementary strain Δ*acoD*/pBBR1MCS-*acoD*, and the mutant strain complemented with an empty vector (Δ*acoD*/pBBR1MCS). All strains were grown with DMSP as the sole carbon source. The error bar represents standard deviation of triplicate experiments. **b**, GC detection of DMS production from DMSP degradation by the wild-type strain D2, the Δ*acoD* mutant, the complemented mutant Δ*acoD*/pBBR1MCS-*acoD*, and the mutant complimented with an empty vector Δ*acoD*/pBBR1MCS. The DMS standard was used as a positive control. **c**, GC detection of DMS production from DMSP lysis catalyzed by the recombinant AcoD. The reaction system without AcoD was used as the control. **d**, HPLC assay of the enzymatic activity of the recombinant AcoD on DMSP. The peak of the unknown product was indicated with a red arrow. The reaction system without AcoD was used as the control. **e**, LC-MS analysis of the unknown product. **f**, LC-MS analysis of the intermediate of AcoD catalysis. The reaction system without AcoD was used as the control.

### The catalytic process of AcoD as an ATP-dependent DMSP lyase

To verify the enzymatic activity of AcoD on DMSP, we cloned the gene *acoD*, expressed it in *Escherichia coli* BL21 (DE3), and purified the recombinant AcoD (Supplementary Fig. 4). Sequence analysis suggests that AcoD is an acetate-CoA ligase, which belongs to the acyl-CoA synthetase (ACD) superfamily and requires CoA and ATP as cofactors for catalysis (***Musfeldt et al., 2002; Mai et al., 1996***). Thus, we added CoA and ATP into the reaction system when measured the enzymatic activity of the recombinant AcoD on DMSP. GC detection showed that the recombinant AcoD could directly act on DMSP to produce DMS (Fig. 3c). Detection of the other products of this reaction using HPLC showed that ADP and an unknown product were generated (Fig. 3d). The chromatographic retention time of the unknown product was the same as that of acryloyl-CoA (***Wang et al., 2017; Cao et al., 2017***), suggesting that this product is likely acryloyl-CoA. This is further supported by liquid chromatography-mass spectrometry (LC-MS) analysis. The molecular weight (MW) of the component in the peak of the unknown product is 822.1317, which is equal to that of acryloyl-CoA (Fig. 3e). Altogether, these results demonstrate that AcoD is a functional ATP-dependent DMSP lyase that can catalyze DMSP degradation to DMS and acryloyl-CoA.

The biochemical results above suggest that AcoD catalyzes a two-step reaction for DMSP catabolism, a CoA ligation reaction and a cleavage reaction. To perform this two-step reaction, there are two alternative pathways: (i), DMSP is first cleaved to form DMS and acrylate, and subsequently CoA is ligated with acrylate (Supplementary Fig. 5a). In this case, the intermediate acrylate is produced. (ii), CoA is ligated with DMSP firstly, and DMSP-CoA is generated. Then, DMSP-CoA is cleaved, producing DMS and acryloyl-CoA (Supplementary Fig. 5b). In this scenario, the intermediate DMSP-CoA is produced. To determine the catalytic process of AcoD, we monitored the presence of acrylate and/or DMSP-CoA in the reaction system via LC-MS. While acrylate was not detectable in the reaction system, a small peak of DMSP-CoA emerged after a 2-min reaction (Fig. 3f), indicating that DMSP-CoA is firstly formed in the catalytic reaction of AcoD, which is then cleaved to generate DMS and acryloyl-CoA.

### The crystal structure and the catalytic mechanism of AcoD

To elucidate the structure basis of AcoD catalysis, we solved the crystal structure of AcoD in complex with ATP by the single-wavelength anomalous dispersion method using a selenomethionine derivative (Se-derivative) (Supplementary Table 3). Although there are four AcoD monomers arranged as a tetramer in an asymmetric unit (Supplementary Fig. 6a), gel filtration analysis indicated that AcoD maintains a dimer in solution (Supplementary Fig. 6b). Each AcoD monomer contains a CoA-binding domain and an ATP-grasp domain (Fig. 4a), with one loop (Gly280-Tyr300) of the CoA-binding domain inserting into the ATP-grasp domain. ATP is bound in AcoD mainly via hydrophilic interactions, including hydrogen bonds and salt bridges (Fig. 4b). The overall structure of AcoD is similar to that of NDP-forming acetyl-CoA synthetase (ACS) ACD1 (***Weiße et al., 2016***) (Supplementary Fig. 7), with a root mean square deviation (RMSD) between these two structures of 4.6 Å. ACD1 consists of separate α- and β-subunits (***Weiße et al., 2016***), which corresponds to the CoA-binding domain and the ATP-grasp domain of AcoD, respectively.

**Fig. 4.**
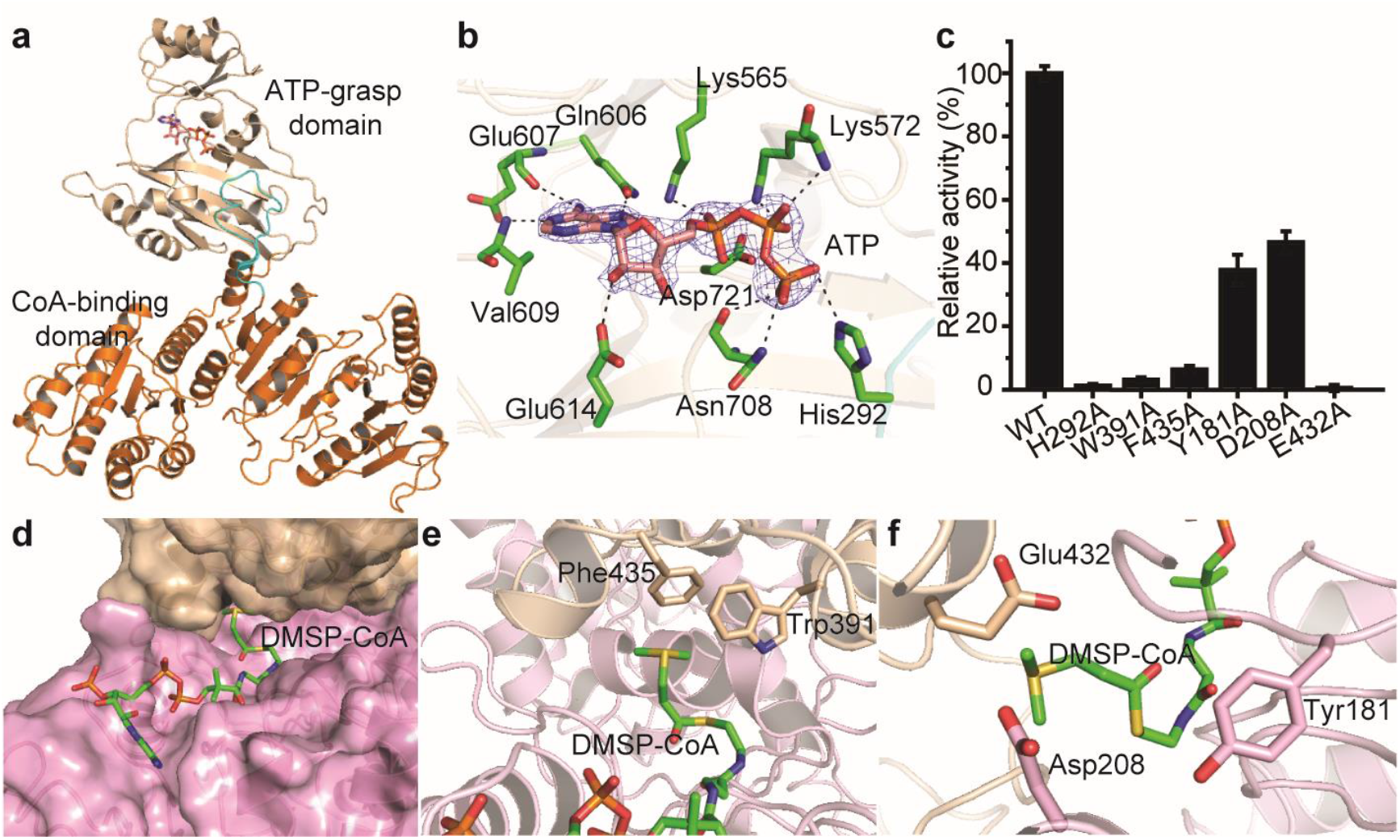
Structural and mutational analyses of AcoD. **a**, The overall structure of AcoD monomer. The AcoD molecule contains a CoA-binding domain (colored in orange) and an ATP-grasp domain (colored in wheat). The loop region from the CoA-binding domain inserting into the ATP-grasp domain is colored in cyan. The ATP molecule is shown as sticks. **b**, Residues of AcoD involved in binding ATP. The 2*F*_*o*_ - *F*_*c*_ densities for ATP are contoured in blue at 2.0σ. Residues of AcoD involved in binding ATP are colored in green. **c**, Enzymatic activities of AcoD and its mutants. The activity of WT AcoD was taken as 100%. **d**, The structure of AcoD docked with DMSP-CoA. DMSP-CoA is shown as sticks. The surfaces of two AcoD monomers are colored in wheat and pink, respectively. **e**, Structural analysis of residues which form cation-π interactions with the sulfonium group of DMSP-CoA. DMSP-CoA and residues Trp391 and Phe435 are shown as sticks. **f**, Structural analysis of the possible catalytic residues for the cleavage of DMSP-CoA. DMSP-CoA and the probable catalytic residues of AcoD are shown as sticks.

Both AcoD and ACS belong to the ACD superfamily, which also contains the well-studied ATP citrate lyases (ACLY) (***Weiße et al., 2016; Verschueren et al., 2019; Hu et al., 2017***). The biochemistry of AcoD catalysis is similar to that of ACLY, which converts citrate to acetyl-CoA and oxaloacetate with ATP and CoA as cofactors (***Verschueren et al., 2019; Hu et al., 2017***). Phylogenetic analysis indicated that AcoD and its homologs form a separate clade from ACS and ACLY, suggesting the divergent evolution of AcoD from ACS and ACLY (Supplementary Fig. 8). The catalytic processes of enzymes in the ACD superfamily involve a conformational change of a “swinging loop” or “phosphohistidine segment”, in which a conserved histidine is phosphorylated (***Weiße et al., 2016; Verschueren et al., 2019; Hu et al., 2017***). Sequence alignment indicated that His292 of AcoD is the conserved histidine residue to be phosphorylated, and Gly280-Tyr300 is likely the “swinging loop” (Supplementary Fig. 9). In the crystal structure of AcoD, His292 from loop Gly280-Tyr300 directly forms a hydrogen bond with the γ-phosphate of ATP (Fig. 4b), suggesting a potential of phosphorylation, which is further supported by mutational analysis. Mutation of His292 to alanine abolished the activity of AcoD (Fig. 4c), indicating the key role of His292 during catalysis. Circular-dichroism (CD) spectroscopy analysis showed that the secondary structure of His292Ala exhibits little deviation from that of wild-type (WT) AcoD (Supplementary Fig. 10), indicating that the enzymatic activity loss was caused by amino acid replacement rather than by structural change. Altogether, these data suggest that His292 is phosphorylated in the catalysis of AcoD on DMSP.

Having solved the crystal structure of AcoD-ATP complex, we next sought to determine the crystal structures of AcoD in complex with CoA and DMSP. However, the diffractions of these crystals are poor and all attempts to solve the structures were failed. Then we docked DMSP and CoA in the structure of AcoD. In the docked structure, the CoA molecule is bound in the CoA-binding domain, while the DMSP molecule is bound in the interface between two AcoD monomers (Supplementary Fig. 11). Because our biochemical results demonstrated that DMSP-CoA is an intermediate of AcoD catalysis (Fig. 3f), we further docked DMSP-CoA in AcoD. DMSP-CoA also locates between two AcoD monomers (Fig. 4d), and two aromatic residues (Trp391 and Phe435) form cation-π interactions with the sulfonium group of DMSP-CoA (Fig. 4e). Mutations of these two residues significantly decreased the enzymatic activities of AcoD (Fig. 4c), suggesting that these residues play important roles in stabilizing the DMSP-CoA intermediate. To cleave DMSP-CoA into DMS and acryloyl-CoA, a catalytic residue is necessary to attack the DMSP moiety. Structure analysis showed that Tyr181, Asp208 and Glu432 are close to the DMSP moiety (Fig. 4f) and may function as the general base. Mutational analysis showed that the mutation of Glu432 to alanine abolished the enzymatic activity of AcoD, while mutants Tyr181Ala and Asp208Ala still maintained ~40% activities (Fig. 4c), indicating that Glu432 is the most probable catalytic residue for the final cleavage of DMSP-CoA. CD spectra of these mutants were indistinguishable from that of WT AcoD (Supplementary Fig. 10), suggesting that the decrease in the enzymatic activities of the mutants are caused by residue replacement rather than structural alteration of the enzyme.

Based on structural and mutational analyses of AcoD, and the reported molecular mechanisms of the ACD superfamily (***Weiße et al., 2016; Verschueren et al., 2019; Hu et al., 2017***), we proposed the molecular mechanism of AcoD catalysis on DMSP (Fig. 5). Firstly, His292 is phosphorylated by ATP, forming phosphohistidine (Fig. 5a), which will be brought to the CoA-binding domain through the conformational change of the swinging loop Gly280-Tyr300. Next, the phosphoryl group is transferred to DMSP to generate DMSP-phosphate (Fig. 5b), which is subsequently attacked by CoA to form DMSP-CoA intermediate (Fig. 5c). The last step is the cleavage of DMSP-CoA initiated by the nucleophilic attack of Glu432 (Fig. 5d). Finally, acryloyl-CoA and DMS are generated (Fig. 5e) and released from the catalytic pocket of AcoD.

**Fig. 5.**
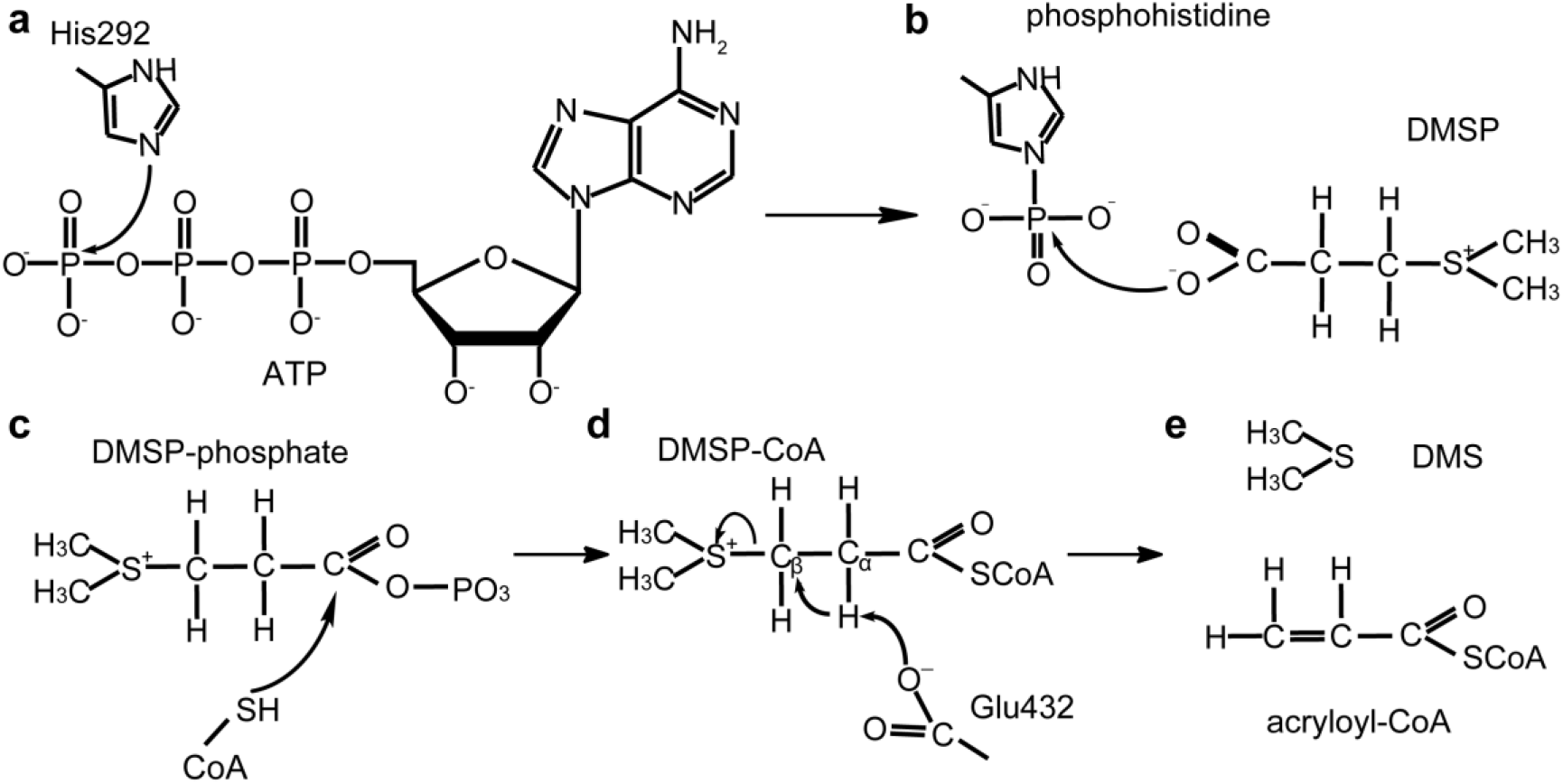
A proposed mechanism for DMSP cleavage to generate DMS and acryloyl-CoA catalyzed by AcoD. **a**, The residue His292 attacks the γ-phosphate of ATP. **b**, The phosphoryl group is transferred from phosphohistidine to the DMSP molecule. **c**, DMSP-phosphate is attacked by CoA. **d**, The residue Glu432 acts as a general base to attack DMSP-CoA. **e**, DMS and acryloyl-CoA are generated.

### A novel ATP DMSP lysis pathway in *Psychrobacter* sp. D2

It has been reported that accumulation of acryloyl-CoA is toxic to bacteria (***Reisch et al., 2013; Wang et al., 2017; Cao et al., 2017; Todd et al., 2012***). Thus, *Psychrobacter* sp. D2 requires an efficient system to metabolize acryloyl-CoA produced from DMSP lysis by AcoD. Genes *1698* and *1699* are located in the downstream of *acoD*, and the transcriptions of *1698* and *1699* are significantly enhanced by the induction of DMSP. Thus, 1698 and 1699 are likely to participate in the metabolism of acryloyl-CoA. However, the recombinant 1698 and 1699 exhibited little enzymatic activity on acryloyl-CoA. In the genome of *Psychrobacter* sp. D2, we also found *acuI* and *acuH* homologs (*2674*, *0105*, *1810*, *1692* and *1695*) (Supplementary Table 4), which may directly act on acryloyl-CoA to produce propionate-CoA or 3-HP-CoA (***Reisch et al., 2013; Wang et al., 2017; Cao et al., 2017; Todd et al., 2012***). To verify their functions, we purified these proteins, and measured their enzymatic activities towards acryloyl-CoA. HPLC analysis showed that the recombinant 0105, an AcuI homolog, could act on acryloyl-CoA to produce propionate-CoA with NADPH as a cofactor (Supplementary Fig. 12).

Based on our genetic and biochemical analyses, the DMSP catabolic pathway in *Psychrobacter* sp. D2 is proposed. In this pathway, DMSP is first converted to DMS and acryloyl-CoA by AcoD, with DMSP-CoA as an intermediate. DMS can diffuse out from cells and acryloyl-CoA is further metabolized to propionate-CoA by 0105 (AcuI). The produced propionate-CoA could participate in the central carbon metabolism to provide carbon and energy for bacterial growth (***Reisch et al., 2011a***). Because this pathway is initiated by AcoD, an ATP-dependent DMSP lyase, we name this novel DMSP catabolic pathway the ATP DMSP lysis pathway (Fig. 1).

### Distribution of the ATP DMSP lysis pathway in bacteria

We next set out to determine the diversity and distribution of AcoD in genome sequenced bacteria. We searched the NCBI Reference Sequence Database using the AcoD sequence of *Psychrobacter* sp. D2 as the query. The data presented in Fig. 6 showed that AcoD homologs are presented in a wide range of bacteria, including Alphaproteobacteria, Betaproteobacteria, Gammaproteobacteria and Firmicutes. Multiple sequence alignment showed the presence of the key residues involved in phosphorylation (His292) and co-ordination of the substrate (e.g. W391), suggesting that these AcoD homologs are potentially functional in DMSP catabolism in these bacteria. Interestingly, AcoD homologs are also found in many metagenome-resolved genomes from a variety of environmental samples, including marine seawater and sediment, soil and engineered microbial systems, suggesting that AcoD-mediated DMSP degradation may be widely spread in the environment. To further validate that these AcoD homologs are indeed *bona fide* DMSP degrading enzymes, we chemically synthesized and characterized two representative sequences (*Sporosarcina* sp. P33; *Psychrobacter* sp. P11G5), both of which showed activities towards DMSP degradation (Supplementary Fig. 13).

**Fig. 6.**
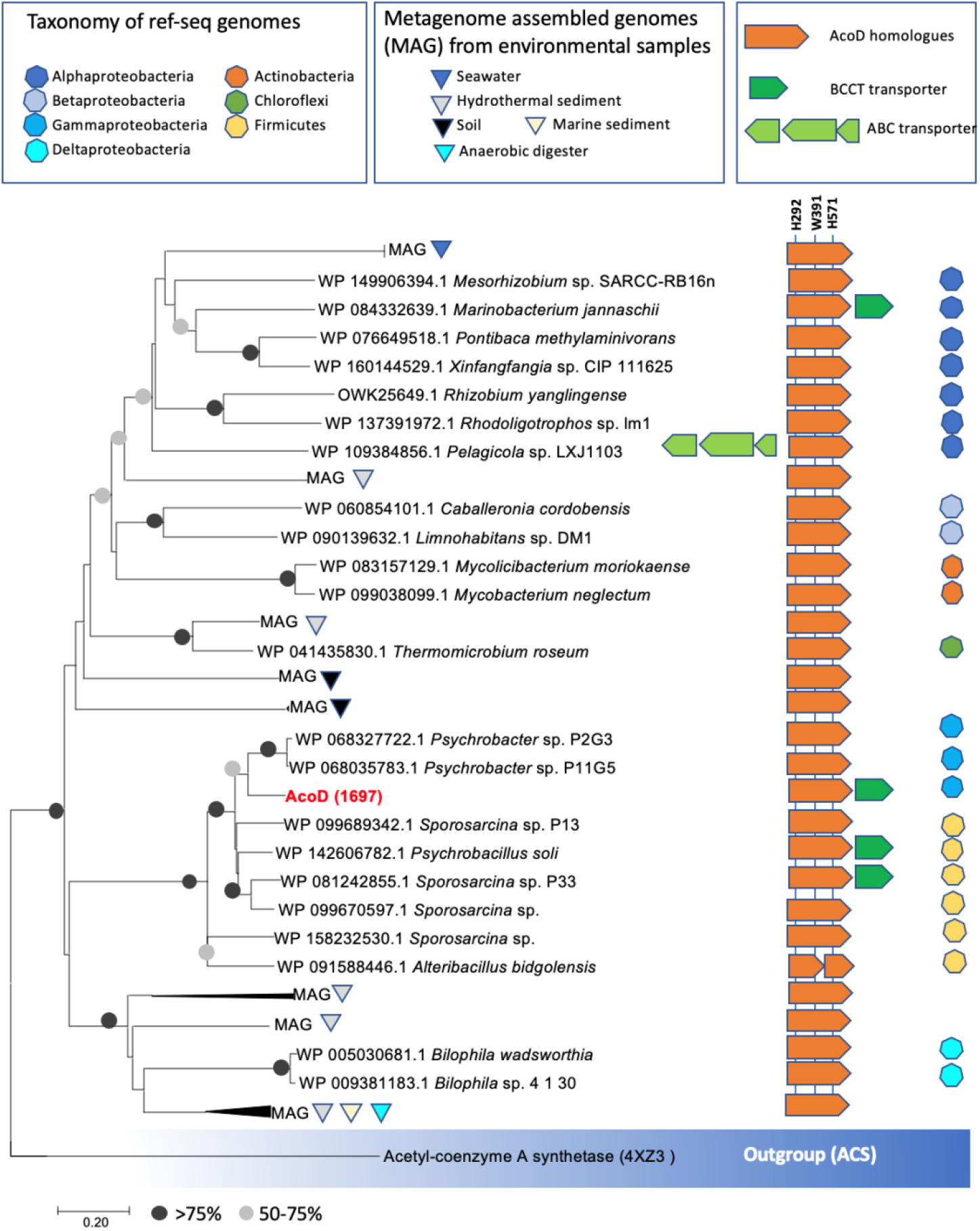
Distribution of AcoD in bacterial genomes and environmental bacteria. The phylogenetic tree was constructed using neighbor-joining method in MEGA7. The acetyl-coenzyme A synthetase (ACS) (***Weiße et al., 2016***) was used as the outgroup. Sequence alignment was inspected for the presence of the key histidine residue (His292) involved in histidine phosphorylation that is known to be important for enzyme activity. A conserved Tyr391 is also found which is involved in cation-pi interaction with DMSP. The BCCT-type or ABC-type transporters for betaine-carnitine-choline-DMSP were found in the neighborhood of AcoD in several genomes.

## Discussion

The cleavage of DMSP to produce DMS is a globally important biogeochemical reaction. Although all known DMSP lyases catalyze this reaction, they belong to very different families, and likely evolved independently (***Bullock et al., 2017***). DddD belongs to the type III acyl CoA transferase family (***Todd et al., 2007***), while DddP belongs to the M24 metallopeptidase family (***Todd et al., 2009***), DddL/Q/W/K/Y belong to the cupin superfamily (***Lei et al., 2018; Li et al., 2017***) and Alma1 belongs to the aspartate racemase superfamily (***Alcolombri et al., 2015***). To the best of our knowledge, AcoD represents the first DMSP lyase of the ACD superfamily.

Among the reported DMSP lyases, only DddD catalyzes a two-step reaction which contains a CoA transfer reaction and a cleavage reaction (***Alcolombri et al., 2014***). It is deduced that DMSP-CoA may be generated in the catalytic process of DddD (***Alcolombri et al., 2014; Curson et al., 2011; Todd et al., 2007***). Despite this, AcoD is fundamentally different from DddD. Firstly, the cofactors of AcoD and DddD are different. ATP and CoA are essential cofactors for the enzymatic activity of AcoD, while for DddD catalysis, acetyl-CoA is used as a CoA donor, and ATP is not needed (***Johnston et al., 2016; Alcolombri et al., 2014***). When CoA was replaced by acetyl-CoA in the reaction system, AcoD failed to catalyze the cleavage of DMSP (Supplementary Fig. 14). Secondly, the products of DddD and AcoD are different. DddD converts DMSP to DMS and 3-HP, whereas AcoD produces DMS and acryloyl-CoA from DMSP. Except for DddD and AcoD, all the other DMSP lyases cleave DMSP to DMS and acrylate. In fact, in the PrpE-AcuI pathway for acrylate metabolism (***Wang et al., 2017***), acryloyl-CoA is an essential intermediate, which is generated from acrylate by PrpE (Fig. 1). Thus, bacteria containing AcoD may not need PrpE homologs to utilize DMSP as a carbon source.

Many marine bacteria, especially roseobacters, are reported to metabolize DMSP via more than one pathway (***Curson et al., 2011; Bullock et al., 2017***). For example, *Ruegeria pomeroyi* DSS-3, one of the type strains of marine *Roseobacter* clade, possesses both the demethylation and the lysis pathway for DMSP metabolism (***Reisch et al., 2013***). Moreover, it contains multiple *ddd* genes (*dddQ*, *dddP* and *dddW*) (***Reisch et al., 2013; Todd et al., 2011)***. DmdA homologs were not identified form the genome of *Psychrobacter* sp. D2, indicating that the demethylation pathway does not exist in strain D2. The fact that the mutant Δ*acoD* could not produce DMS from DMSP and was unable to grow on DMSP as the sole carbon source suggests that *Psychrobacter* sp. D2 only possesses the ATP DMSP lysis pathway for DMSP degradation. Why some bacteria have evolved multiple DMSP utilization pathways and some bacteria only possess one pathway awaits further investigation.

It has been reported that Gram-positive actinobacteria can make DMS from DMSP (***Liu et al., 2018***). However, the responsible enzymes have not been idenfied yet. Here, for the fisrt time, we identified that AcoD homolog is a functional DMSP lyase in *Sporosarcina* sp. P33, a Gram-positive bacterium which belongs to Firmicutes. The wide distribution of AcoD in many bacterial lineages points to its horizontal gene transfer (HGT) event.

## Conclusion

DMSP is widespread in nature and cleavage of DMSP produces DMS, an important mediator in the global sulfur cycle. In this study, we reported the identification of a novel ATP DMSP lysis pathway from marine bacteria. The first enzyme involved in this pathway is an ATP-dependent DMSP lyase AcoD, which belongs to the ACD superfamily. AcoD catalyzes the conversion of DMSP to DMS and acryloyl-CoA, with CoA and ATP as cofactors. AcoD homologs are widely distributed in a range of bacteria. This study offers new insights into how bacteria cleave DMSP to generate the climatically important gas DMS using a novel ATP DMSP lysis pathway.

## Methods

### Bacterial strains, plasmids and growth conditions

Strains and plasmids used in this study are shown in Supplementary Table 5. *Psychrobacter* sp. D2 and other isolates were cultured in the marine broth 2216 medium at 15-25°C. The *E. coli* strains DH5α and BL21(DE3) were grown in the Lysogeny Broth (LB) medium at 37°C. Diaminopimelic acid (0.3 mM) was added to culture the *E. coli* WM3064 strain.

### Isolation of bacterial strains from Antarctic samples

A total of five samples were collected from the Great Wall Station of Antarctica during the Chinese Antarctic Great Wall Station Expedition in January, 2017. Information of samples is shown in Supplementary Fig. 1a and Supplementary Table 1. Algae and sediments were collected using a grab sampler and stored in airtight sterile plastic bags at 4°C. Seawater samples were filtered through polycarbonate membranes with 0.22 μm pores (Millipore Co., United States). The filtered membranes were stored in sterile tubes (Corning Inc., United States) at 4°C. All samples were transferred into a 50 ml flask containing 20 ml 3% (w/v) seasalt solution (SS) and shaken at 100 rpm at 15°C for 2 h. The suspension obtained was subsequently diluted to 10^−6^ with sterile SS. An aliquot (200 μl) of each dilution was spread on the basal medium (Supplementary Table 6) plates with 5 mM DMSP as the sole carbon source. The plates were then incubated at 15°C in the dark for 2-3 weeks. Colonies with different appearances were picked up and were further purified by streaking on the marine 2216 agar plates for at least three passages. The abilities of the colonies for DMSP catabolism were verified in a liquid basal medium with DMSP (5 mM) as the sole carbon source. The isolated were stored at −80°C in the marine broth 2216 medium containing 20% (v/v) glycerol.

### Sequence analysis of bacterial 16S rRNA genes

Genomic DNA of the isolates was extracted using a bacterial genomic DNA isolation kit (BioTeke Corporation, China) according to the manufacturer’s instructions. The 16S rRNA genes of these strains were amplified using the primers 27F/1492R (Supplementary Table 7) and sequenced to determine their taxonomy. Pairwise similarity values for the 16S rRNA gene of the cultivated strains were calculated through the EzBiocloud server (http://www.ezbiocloud.net/) (***Yoon et al., 2017***).

### Bacterial growth assay with DMSP as the sole carbon source

Cells were grown in the marine broth 2216 medium, harvested after incubation at 15°C for 24 h, and then washed three times with sterile SS. The washed cells were diluted to the same density of OD_600_ ≈ 2.0, and then 1% (v/v) cells were inoculated into the basal medium with DMSP (5 mM) as the sole carbon source. The bacteria were cultured in the dark at 15°C. The growth of the bacteria was measured by detecting the OD_600_ of the cultures at different time points using a spectrophotometer V-550 (Jasco Corporation, Japan).

### Assays for DMS production of bacteria strains

To measure the production of DMS, cells were first cultured overnight in the marine broth 2216 medium. Then a final concentration of 5 mM DMSP was added into the medium. After further incubation for 4 h, the cultures were diluted to 10^−3^ with SS and an aliquot (1 ml) of this dilution was added into a glass vial. DMS was diffused from the liquid with a stream of nitrogen and analyzed on a gas chromatograph (GC-2030, Shimadzu, Japan) equipped with a flame photometric detector (***Liu et al., 2018***). The DMS standard (Sigma, America) was used as a positive control.

### Transcriptome sequencing of *Psychrobacter* sp. D2

Cells of strain D2 were cultured in the marine broth 2216 medium at 180 rpm at 15°C for 24 h. The cells were collected and washed three times with sterile SS, and then cultured in sterile SS at 180 rpm at 15°C for 24 h. Subsequently, the cells were washed twice with sterile SS, and incubated at 4°C for 24 h. After incubation, the cells were harvested and resuspended in sterile SS, which were used as the resting cells. The resting cells were inoculated into the basal medium with DMSP (5 mM) as the sole carbon source, and incubated at 180 rpm at 15°C. When the OD_600_ of the cultures reached 0.3, the cells were harvested. The resting cells and those cultured in the basal medium with sodium pyruvate (5 mM) as the sole carbon source were set up as controls. Total RNA was extracted using a RNeasy Mini Kit (QIAGEN, America) according to the manufacturer’s protocol. After validating the quality, RNA samples were sent to BGI Tech Solutions Co., Ltd (China) for transcriptome sequencing and subsequent bioinformatic analysis.

### Real-Time qPCR analysis

Cells of *Psychrobacter* sp. D2 were cultured in the marine broth 2216 medium at 180 rpm at 15°C to an OD_600_ of 0.8. Then, cells were induced by 5 mM DMSP, and the control group without DMSP was also set up. After 20 min’s induction, total RNA was extracted using a RNeasy Mini Kit (Qiagen, Germany) according to the manufacturer’s instructions. Genomic DNA was removed using gDNA Eraser (Takara, Japan) and cDNA was synthesized using a PrimeScript™ RT reagent Kit. The qPCR was performed on the Light Cycler II 480 System (Roche, Switzerland) using a SYBR® Premix Ex Taq™ (TaKaRa, Japan). Relative expression levels of target genes were calculated using the LightCycler®480 software with the “Advanced Relative Quantification” method. The *recA* gene was used as an internal reference gene. The primers used in this study are shown in Supplementary Table 7.

### Genetic manipulations of *Psychrobacter* sp. D2

Deletion of the *acoD* gene was performed via pK18*mobsacB-*Ery-based homologue recombination (***Wang et al., 2015***). The upstream and downstream homologous sequences of the *acoD* gene were amplified with primer sets *acoD*-UP-F/*acoD*-UP-R and *acoD*-Down-F/*acoD*-Down-R, respectively. Next, the PCR fragments were inserted to the vector pK18*mobsacB-*Ery with *Hind*III/*BamH*I as the restriction sites to generate pK18Ery-*acoD*, which was transferred into *E. coli* WM3064. The plasmid pK18Ery-*acoD* was then mobilized into *Psychrobacter* sp. D2 by intergeneric conjugation with *E. coli* WM3064. To select for colonies in which the pK18Ery-*acoD* had integrated into the *Psychrobacter* sp. D2 genome by a single crossover event, cells were plated on the marine 2216 agar plates containing erythromycin (25 μg/ml). Subsequently, the resultant mutant was cultured in the marine broth 2216 medium and plated on the marine 2216 agar plates containing 10% (w/v) sucrose to select for colonies in which the second recombination event occurred. Single colonies appeared on the plates were streaked on the marine 2216 agar plates containing erythromycin (25 μg/ml), and colonies sensitive to erythromycin were further validated to be the *acoD* gene deletion mutants by PCR with primer pairs of *acoD*-1000-F/*acoD*-1000-R and *acoD*-300Up-F/*acoD*-700Down-R.

For complementation of the Δ*acoD* mutant, the *acoD* gene with its native promoter was amplified using the primers set *acoD*-pBBR1-PF/*acoD*-pBBR1-PR. The PCR fragment was digested with *Kpn*I and *Xho*I, and then inserted into the vector pBBR1MCS to generate pBBR1MCS-*acoD*. This plasmid was then transformed into *E. coli* WM3064, and mobilized into the Δ*acoD* mutant by intergeneric conjugation. After mating, the cells were plated on the marine 2216 agar plates containing kanamycin (80 μg/ml) to select for the complemented mutant. The empty vector pBBR1MCS was mobilized into the Δ*acoD* mutant using the same protocol. Colony PCR was used to confirm the presence of the transferred plasmid. The strains, plasmids and primers used in this study are shown in Supplementary Table 5 and Supplementary Table 7.

### Gene cloning, point mutation and protein expression and purification

The 2247 bp full-length *acoD* gene was amplified from the genome of *Psychrobacter* sp. D2 by PCR using *FastPfu* DNA polymerase (TransGen Biotech, China). The amplified gene was then inserted to the *Nde*I/*Xho*I restriction sites of the pET-22b vector (Novagen, Germany) with a C-terminal His tag. All of the point mutations in AcoD were introduced using the PCR-based method and verified by DNA sequencing. The AcoD protein and its mutants were expressed in *E. coli* BL21 (DE3). The cells were cultured in the LB medium with 0.1 mg/ml ampicillin at 37°C to an OD_600_ of 0.8-1.0 and then induced at 18°C for 16 h with 0.5 mM isopropyl-β-D-thiogalactopyranoside (IPTG). After induction, cells were collected by centrifugation, resuspended in the lysis buffer (50 mM Tris-HCl, 100 mM NaCl, 0.5% glycerol, pH 8.0), and lysed by pressure crusher. The proteins were first purified by affinity chromatography on a Ni^2+^-NTA column (GE healthcare, America), and then fractionated by anion exchange chromatography on a Source 15Q column (GE healthcare, America) and gel filtration on a Superdex G200 column (GE healthcare, America). The Se-derivative of AcoD was overexpressed in *E. coli* BL21 (DE3) under 0.5 mM IPTG induction in the M9 minimal medium supplemented with selenomethionine, lysine, valine, threonine, leucine, isoleucine and phenylalanine. The recombinant Se-derivative was purified using the aforementioned protocol for the wild-type AcoD.

### Enzyme assay and product identification

For the enzymatic activity assay of the AcoD protein, the purified AcoD protein (at a final concentration of 0.1 mM) was incubated with 1 mM DMSP, 1 mM CoA, 1 mM ATP, 2 mM MgCl_2_ and 100 mM Tris-HCl (pH 8.0). The reaction was performed at 37°C for 0.5 h, and terminated by adding 10% (v/v) hydrochloric acid. The control groups had the same reaction system except that the AcoD protein was not added. DMS was detected by GC as described above. Products of acryloyl-CoA and DMSP-CoA were analyzed using LC-MS. Components of the reaction system were separated on a reversed-phase SunFire C_18_ column (Waters, Ireland) connected to a high performance liquid chromatography (HPLC) system (Dionex, United States). The samples were eluted with a linear gradient of 1-20% (v/v) acetonitrile in 50 mM ammonium acetate (pH 5.5) over 24 min. The HPLC system was coupled to an impact HD mass spectrometer (Bruker, Germany) for *m/z* determination.

### Crystallization and data collection

The purified AcoD protein was concentrated to ~ 8 mg/ml in 10 mM Tris–HCl (pH 8.0) and 100 mM NaCl. The AcoD protein was mixed with ATP (1 mM), and the mixtures were incubated at 0°C for 1 h. Initial crystallization trials for AcoD/ATP complex were performed at 18°C using the sitting-drop vapor diffusion method. Diffraction-quality crystals of AcoD/ATP complex were obtained in hanging drops containing 0.1 M lithium sulfate monohydrate, 0.1 M sodium citrate tribasic dihydrate (pH 5.5) and 20% (w/v) polyethylene glycol (PEG) 1000 at 18°C after 2-week incubation. Crystals of the AcoD Se-derivative were obtained in hanging drops containing 0.1 M HEPES (pH 7.5), 10% PEG 6000 and 5% (v/v) (+/−)-2-Methyl-2,4-pentanediol at 18°C after 2-week incubation. X-ray diffraction data were collected on the BL18U1 and BL19U1 beamlines at the Shanghai Synchrotron Radiation Facility. The initial diffraction data sets were processed using the HKL3000 program with its default settings (***Minor et al., 2006***).

### Structure determination and refinement

The crystals of AcoD/ATP complex belong to the C2 space group, and Se-derivative of AcoD belong to the *P*2_1_2_1_2_1_ space group. The structure of AcoD Se-derivative was determined by single-wavelength anomalous dispersion phasing. The crystal structure of AcoD/ATP complex was determined by molecular replacement using the CCP4 program Phaser (***Winn et al., 2011***) with the structure of AcoD Se-derivative as the search model. The refinements of these structures were performed using Coot (***Emsley et al., 2010***) and *Phenix* (***Adams et al., 2010***). All structure figures were processed using the program PyMOL (http://www.pymol.org/).

### Circular dichroism (CD) spectroscopy

CD spectra for WT AcoD and its mutants were carried out in a 0.1 cm-path length cell on a JASCO J-1500 Spectrometer (Japan). All proteins were adjusted to a final concentration of 0.2 mg/ml in 10 mM Tris-HCl (pH 8.0) and 100 mM NaCl. Spectra were recorded from 250 to 200 nm at a scan speed of 200 nm/min.

### Molecular docking simulations

The structure of the AcoD/ATP complex containing a pair of subunits, α & β was loaded and energy minimised in Flare (v3.0, Cresset) involving 11248 moving heavy atoms (Chain A: 5312, Chain B: 5312, Chain G: 10 and Chain S Water: 614). The molecule minimized with 2000 iterations using a gradient of 0.657 kcal/A. The minimised structure had an RMSD 0.82Å relative to the starting structure and a decrease in starting energy from 134999.58 kcal/mol to a final energy of 6888.60 kcal/mol. The DMSP, CoA and DMSP-CoA molecules were drawn in MarvinSketch (v19.10.0, 2019, ChemAxon for Mac) and exported as a Mol SDF format. The molecules were imported into Flare and docked into the proposed CoA/DMSP binding site using the software’s default docking parameters for intensive pose searching and scoring.

### Identification of AcoD homologs in bacteria and phylogenetic analysis

AcoD (1697) of *Psychrobacter* sp. D2 was used as the query sequence to search for homologs in genome-sequenced bacteria in the NCBI Reference Sequence Database (RefSeq, https://www.ncbi.nlm.nih.gov/refseq/) using BLastP with a stringent setting with an e-value cut-off < −80, sequence coverage >70% and percentage identity >50%. These high stringency settings are necessary to exclude other acetyl-CoA synthetase family proteins (ACS) which are unlikely to be involved in DMSP catabolism due to the lack of the conserved histidine 292 or the tryptophan 391 (Fig 6). To confirm the activity of AcoD homologs from genome-sequenced bacteria, two sequences (*Sporosarcina* sp. P33; *Psychrobacter* sp. P11G5) were chemically-synthesized and their enzyme activity for DMSP degradation was confirmed experimentally (Supplementary Fig. 13). A profile hidden Markov model was produced for AcoD using these three activity-validated AcoD using HMMER (***Eddy, 1998***), which was used to search for AcoD homologs in the NCBI non-redundant protein sequences (nr) using the same cut-off settings and AcoD homologs obtained from metagenome-assembled genomes (MAGs) were obtained for sequence alignment and phylogenetic analysis. These AcoD homolog sequences were then aligned using MEGA 7 (***Kumar et al., 2016***) and the phylogenetic tree was constructed using the neighbour-joining method with 500 bootstraps. The characterized ACS ACD1 (***Weiße et al., 2016***) was used as the outgroup.

### Data and materials availability

The draft genome sequences of *Psychrobacter* sp. D2 have been deposited in the National Center for Biotechnology Information (NCBI) Genome database under accession number JACDXZ000000000. All the RNA-seq read data have been deposited in NCBI’s sequence read archive (SRA) under project accession number PRJNA646786. The structure of AcoD/ATP complex has been deposited in the PDB under the accession code 7CM9.

## Acknowledgments

We thank the staffs from BL18U1 & BL19U1 beamlines of National Facility for Protein Sciences Shanghai (NFPS) and Shanghai Synchrotron Radiation Facility, for assistance during data collection. We thank Caiyun Sun and Jingyao Qu from State Key laboratory of Microbial Technology of Shandong University for their help in HPLC and LC-MS.

## Author contributions

C.-Y.L. and X.-J.W. performed the majority of the experiments and data interpretation. Y.C. and M.Q. performed docking analysis. B.R. and Y.C. performed phylogeny analysis. Q.S., S.Z., P.W., X.S. and C.G. helped in experiments. X.-L.C. directed the study. C.-Y.L., X.-J.W. and X.-L.C. wrote the manuscript. Y.C. did critical revision of the manuscript for important intellectual content. Y.-Z.Z. designed the study.

## Supplementary Materials

**Supplementary Figure 1.**
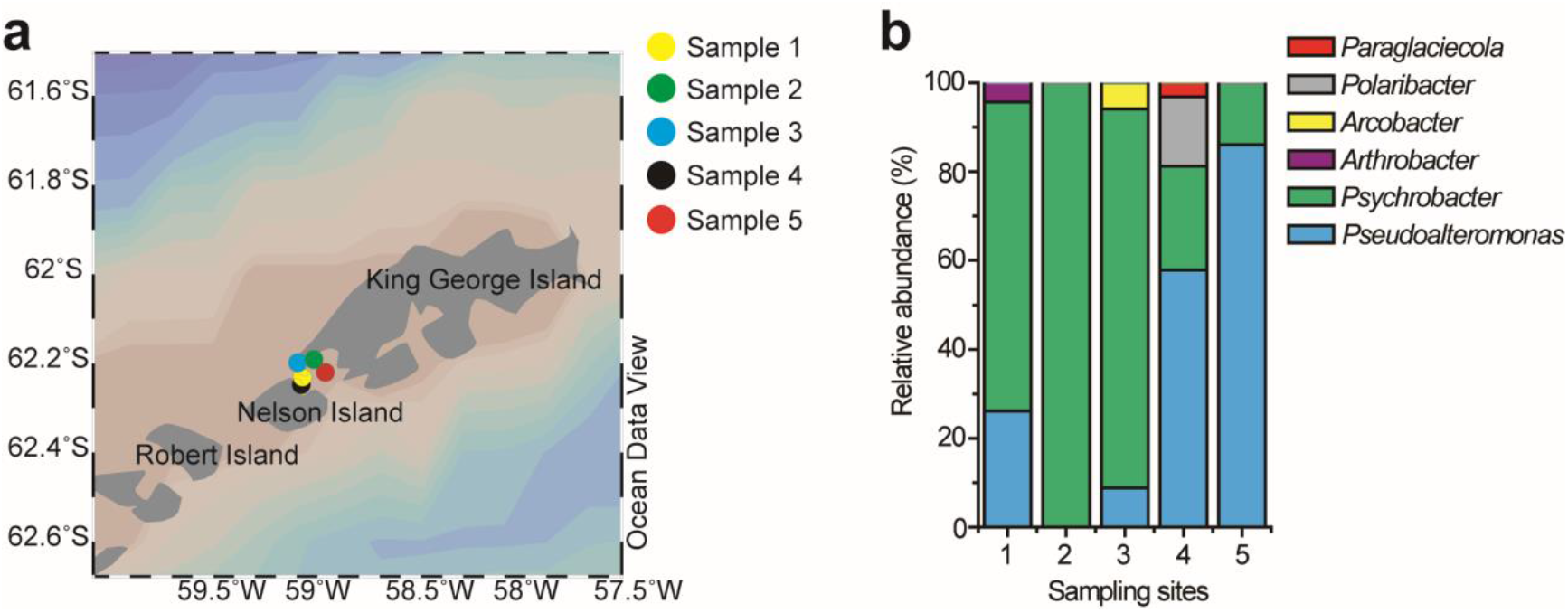
Locations of the sampling sites and the relative abundance of DMSP-catabolizing bacteria isolated from the samples. **a**, Locations of the sampling sites in the Antarctic. Stations were plotted using Ocean Data View (***Schlitzer, 2002***). **b**, The relative abundance of DMSP-catabolizing bacteria isolated from the Antarctic samples. The detail information of the samples is shown in Supplementary Table 1.

**Supplementary Figure 2.**
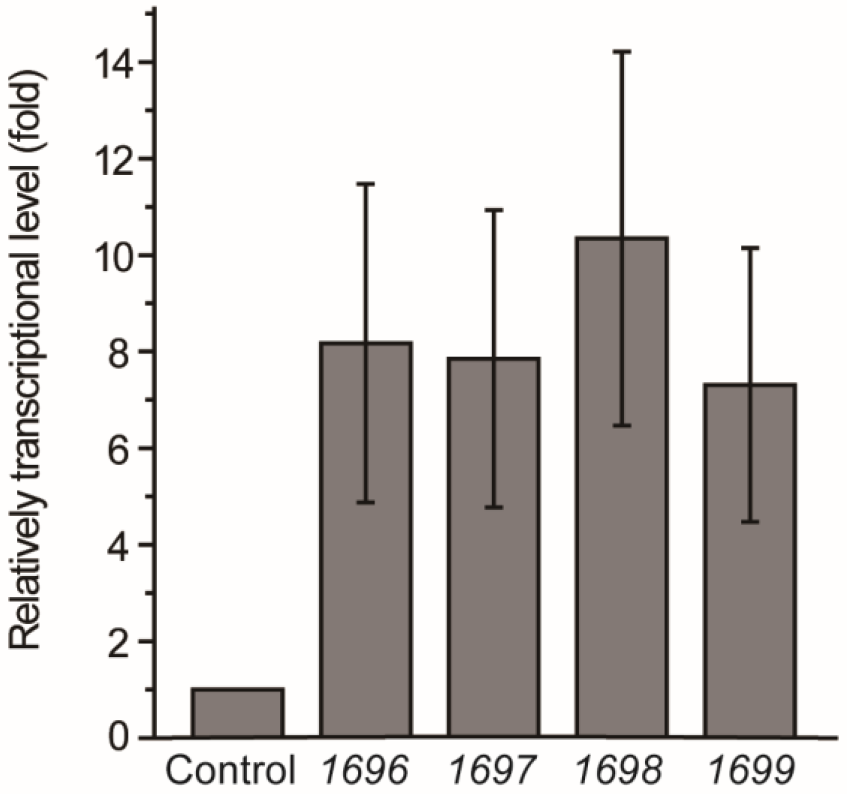
RT-qPCR assay of the transcriptions of the genes *1696*, *1697*, *1698* and *1699* in *Psychrobacter* sp. D2 in response to DMSP in the marine broth 2216 medium. The bacterium cultured without DMSP in the same medium was used as the control. The *recA* gene was used as an internal reference (control). The error bar represents standard deviation of triplicate experiments.

**Supplementary Figure 3.**
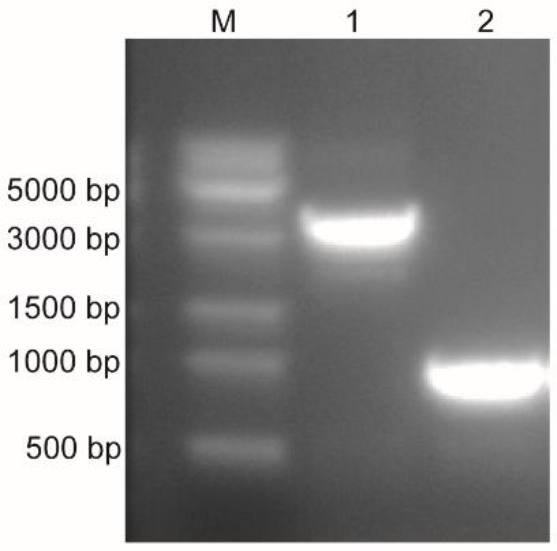
Confirmation of the deletion of the *acoD* gene from *Psychrobacter* sp. D2. Lane M, DNA marker; Lane 1, Wild-type *Psychrobacter* sp. D2; Lane 2, the Δ*acoD* mutant. The Δ*acoD* mutant generated a 1000 bp PCR product using the *acoD*-1000-F/*acoD*-1000-R primer set, while the product length was 3247 bp for the wild-type strain.

**Supplementary Figure 4.**
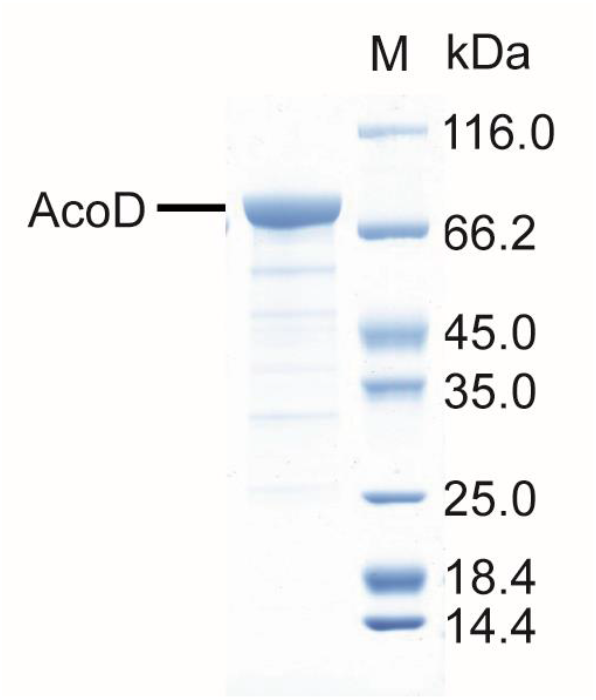
SDS-PAGE analysis of the recombinant AcoD. The predicted molecular mass of the recombinant AcoD is 81.62 kDa using the compute MW tool (***Gasteiger et al., 2005***).

**Supplementary Figure 5.**
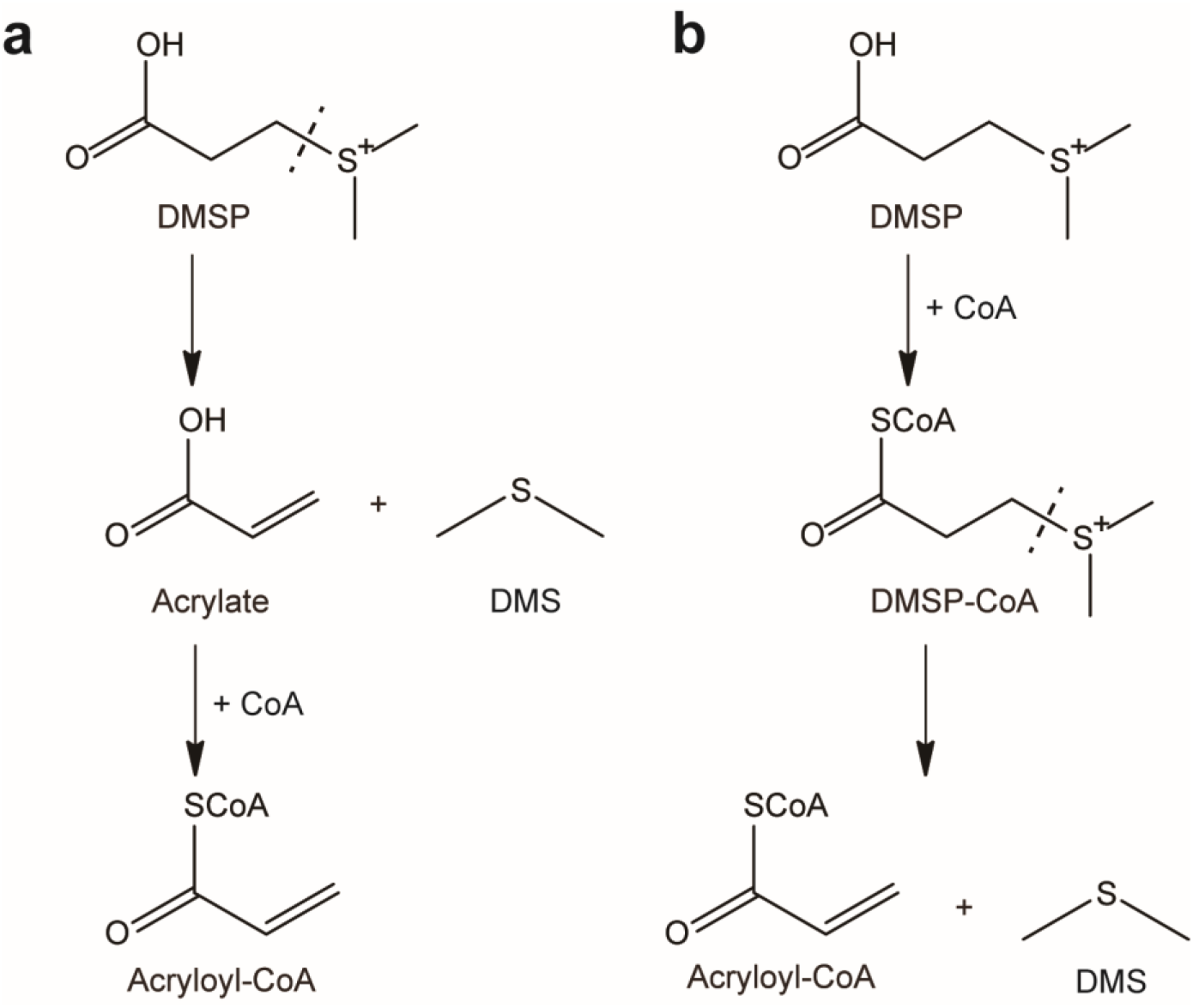
Two alternative mechanisms for DMSP degradation catalyzed by AcoD. **a**, DMSP is cleaved to DMS and acrylate firstly. Subsequently, CoA is ligated to acrylate producing acryloyl-CoA. **b**, DMSP-CoA is generated firstly, which is then cleaved to DMS and acryloyl-CoA.

**Supplementary Figure 6.**
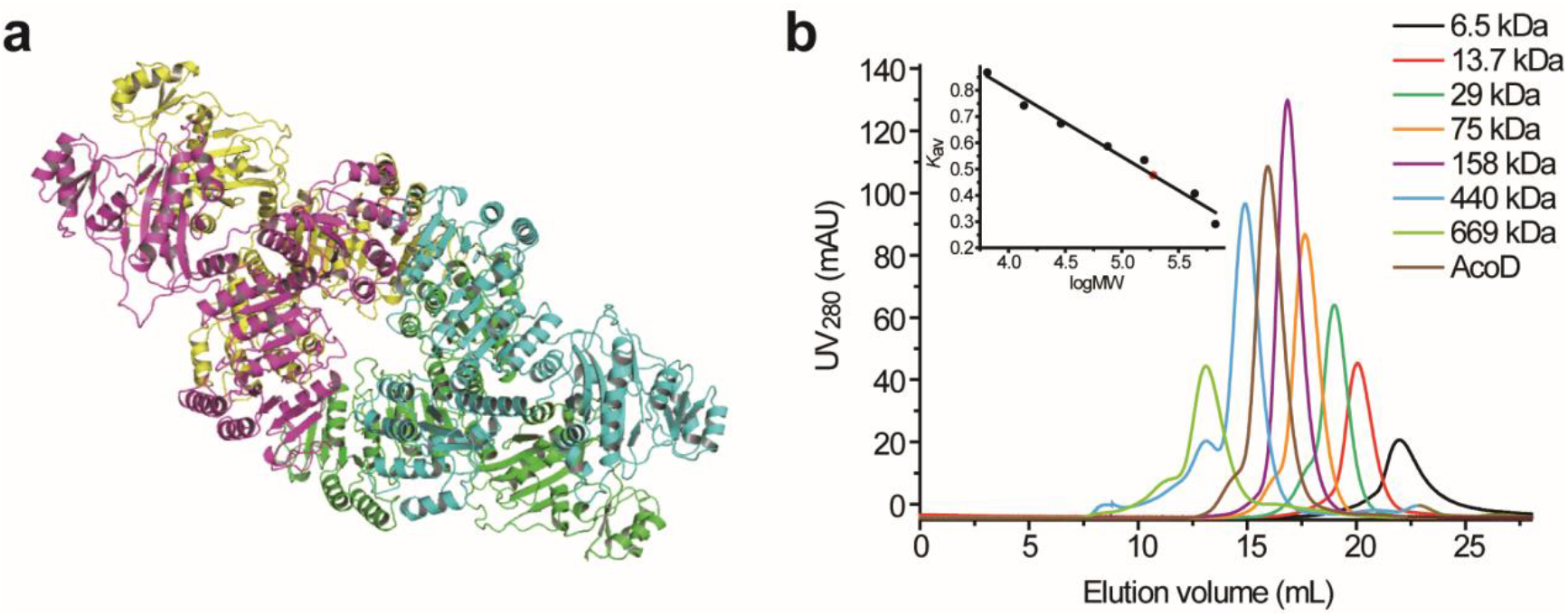
Structural and gel filtration analysis of AcoD state of aggregation. **a**, The overall structure of AcoD tetramer. Different monomers are displayed in different colors. **b**, Gel filtration analysis of AcoD. Inset, semilog plot of the molecular mass of all standards used versus their *K*av values (black circles). The red spot indicates the position of the *K*av value of AcoD interpolated in the regression line. AcoD monomer has a molecular mass of 81.62 kDa.

**Supplementary Figure 7.**
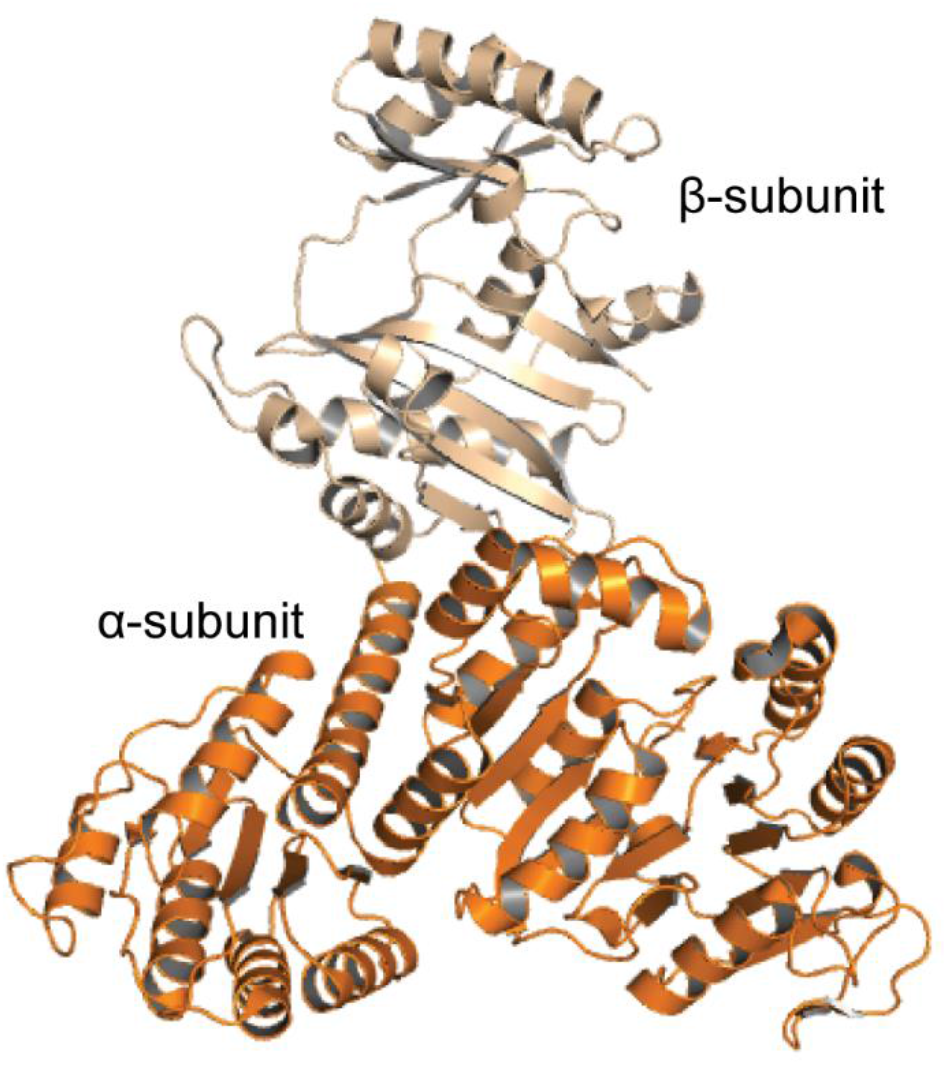
The overall structure of ACD1. The α-subunit and the β-subunit of ACD1 (PDB code: 4xym) are colored in orange and wheat, respectively.

**Supplementary Figure 8.**
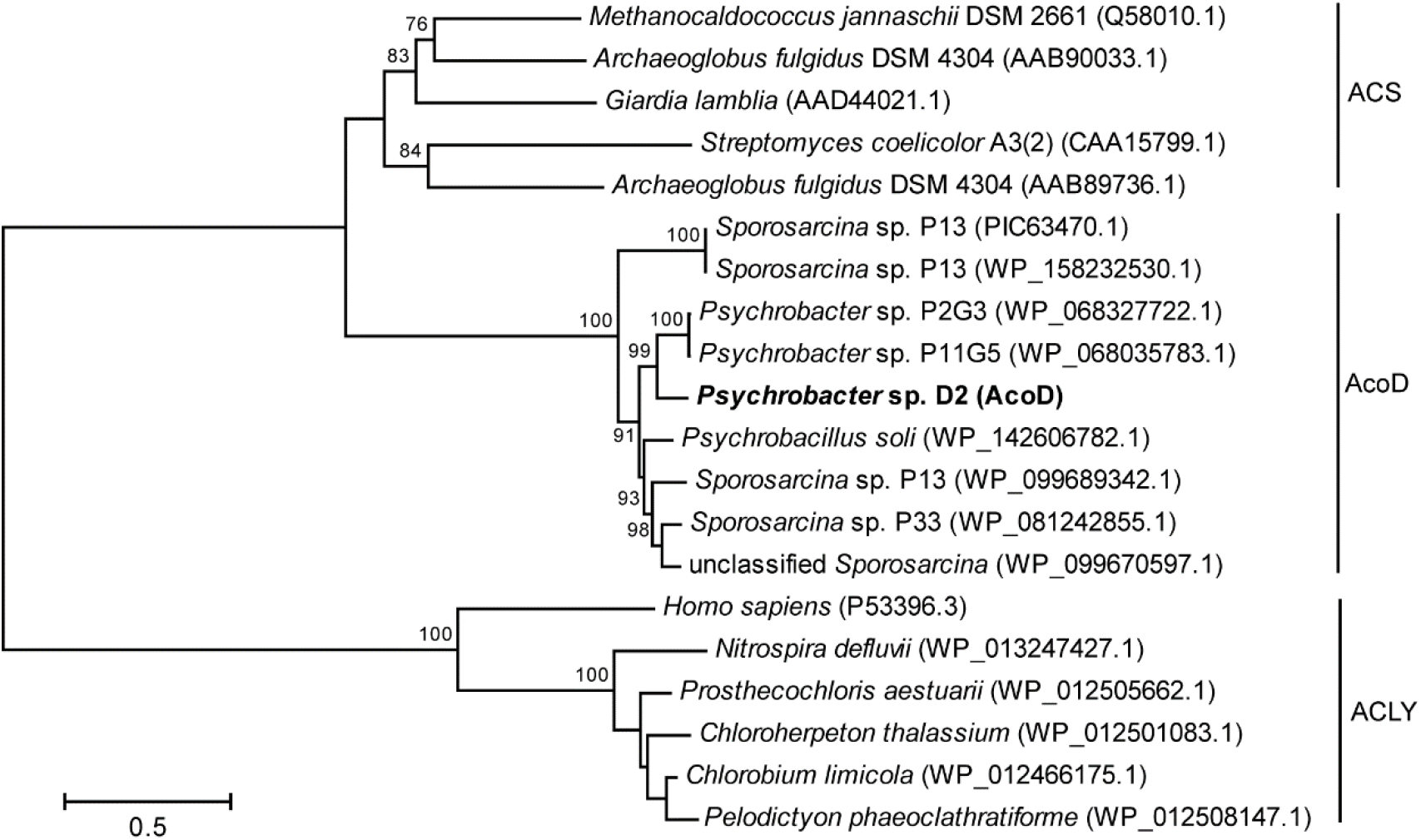
The phylogenetic tree of AcoD and its close homologs, acetyl-CoA synthetases (ACS) and ATP-citrate lyases (ACLY). Bootstrap values of >75 were shown. The scale bar indicates evolutionary distance. Phylogenetic tree was built using neighbor-joining method with 1000 bootstraps.

**Supplementary Figure 9.**
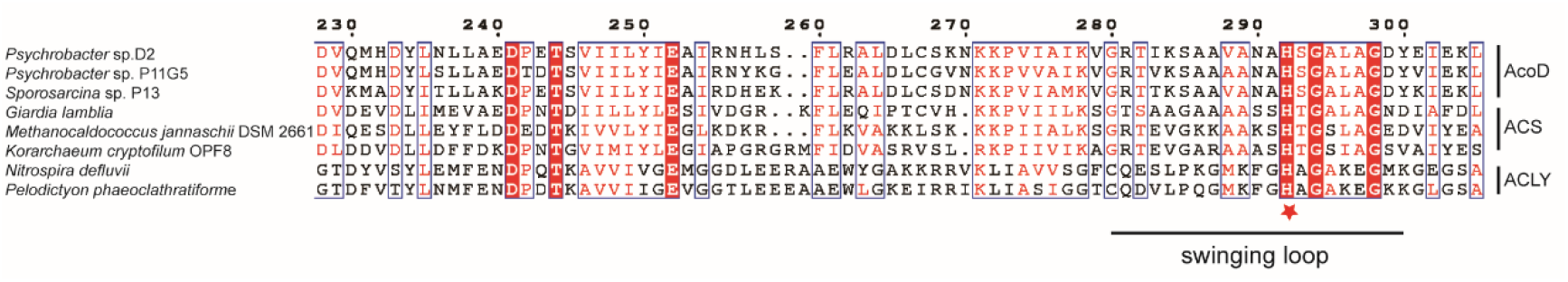
Sequence alignment of AcoD homologs, acetyl-CoA synthetases (ACS) and ATP-citrate lyases (ACLY). The conserved histidine residue is marked with a red star. The swinging loop of AcoD (Gly280-Tyr300) is indicated, which corresponds to the swinging loop reported in acetyl-CoA synthetase ACD1 (Gly242-Val262) (***Weiße et al., 2016***).

**Supplementary Figure 10.**
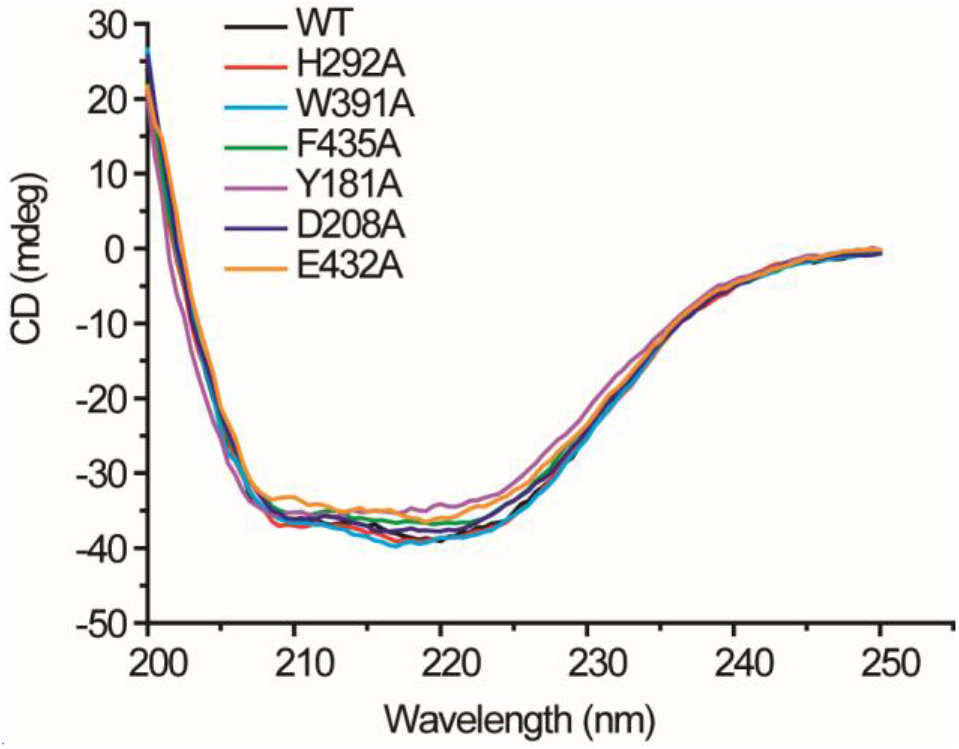
CD spectra of WT AcoD and its mutants.

**Supplementary Figure 11.**
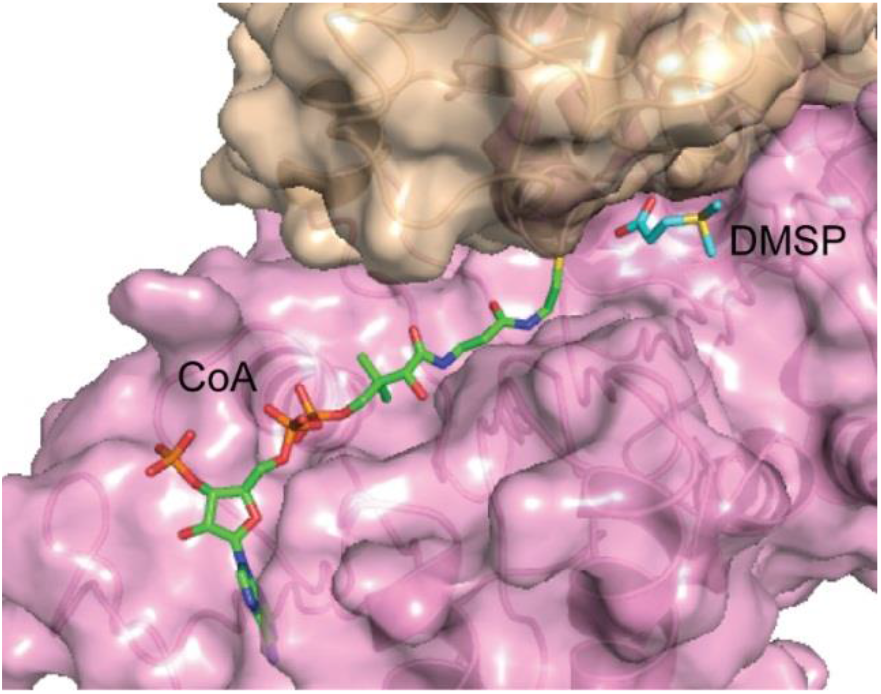
The structure of AcoD docked with DMSP and CoA. DMSP and CoA molecules are shown as sticks. The surfaces of two AcoD monomers are colored in wheat and pink, respectively.

**Supplementary Figure 12.**
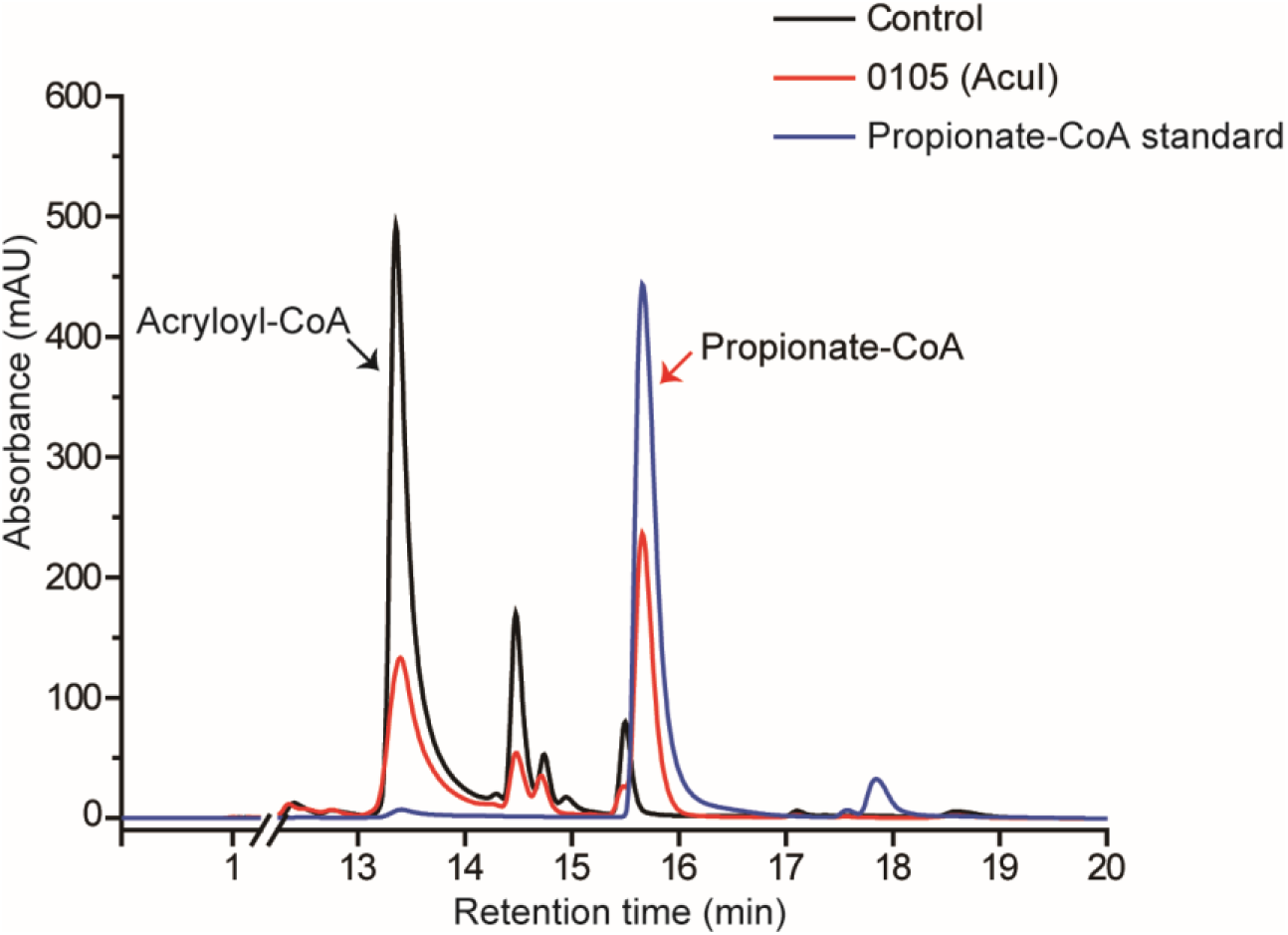
HPLC assay of the enzymatic activity of 0105 protein on acryloyl-CoA. The peak of acryloyl-CoA was indicated with black arrow and the peak of propionate-CoA was indicated with red arrow. The reaction system without 0105 protein was used as the control.

**Supplementary Figure 13.**
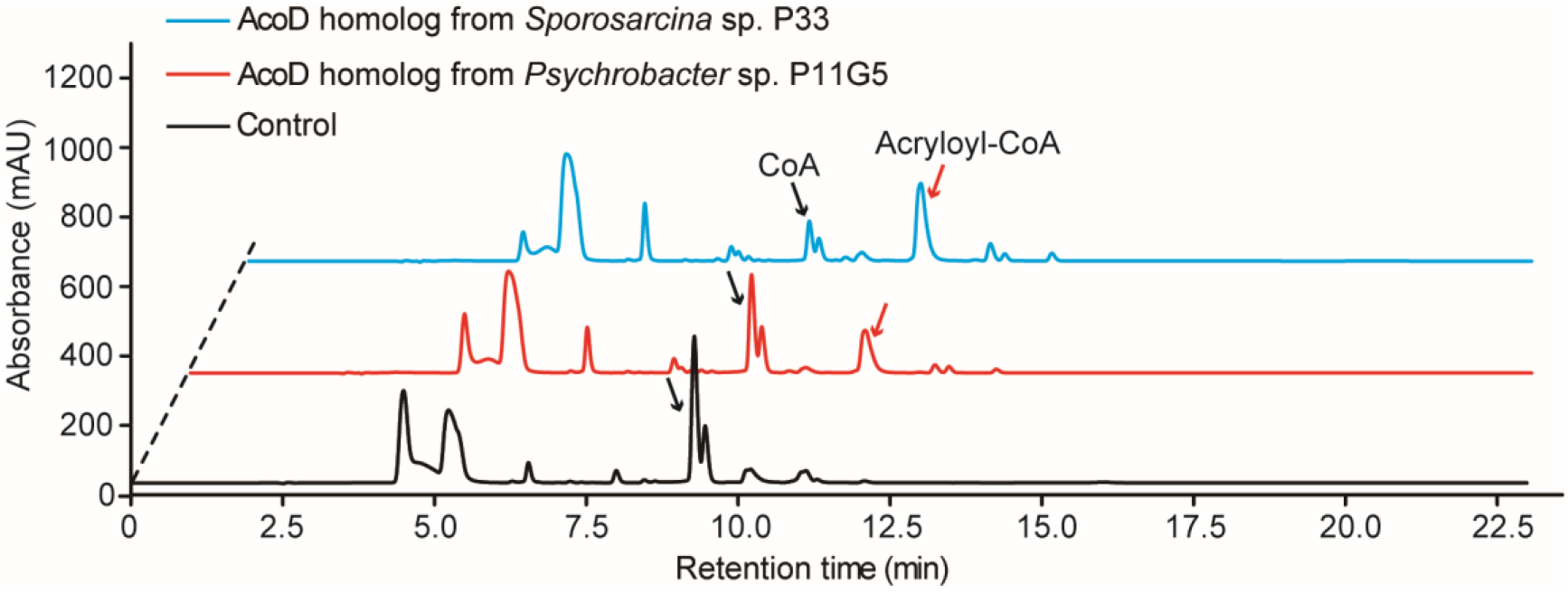
HPLC assay of the enzymatic activity of AcoD homologs on DMSP. The peaks of acryloyl-CoA were indicated with red arrows and the peaks of CoA were indicated with black arrows. The reaction system without AcoD was used as the control.

**Supplementary Figure 14.**
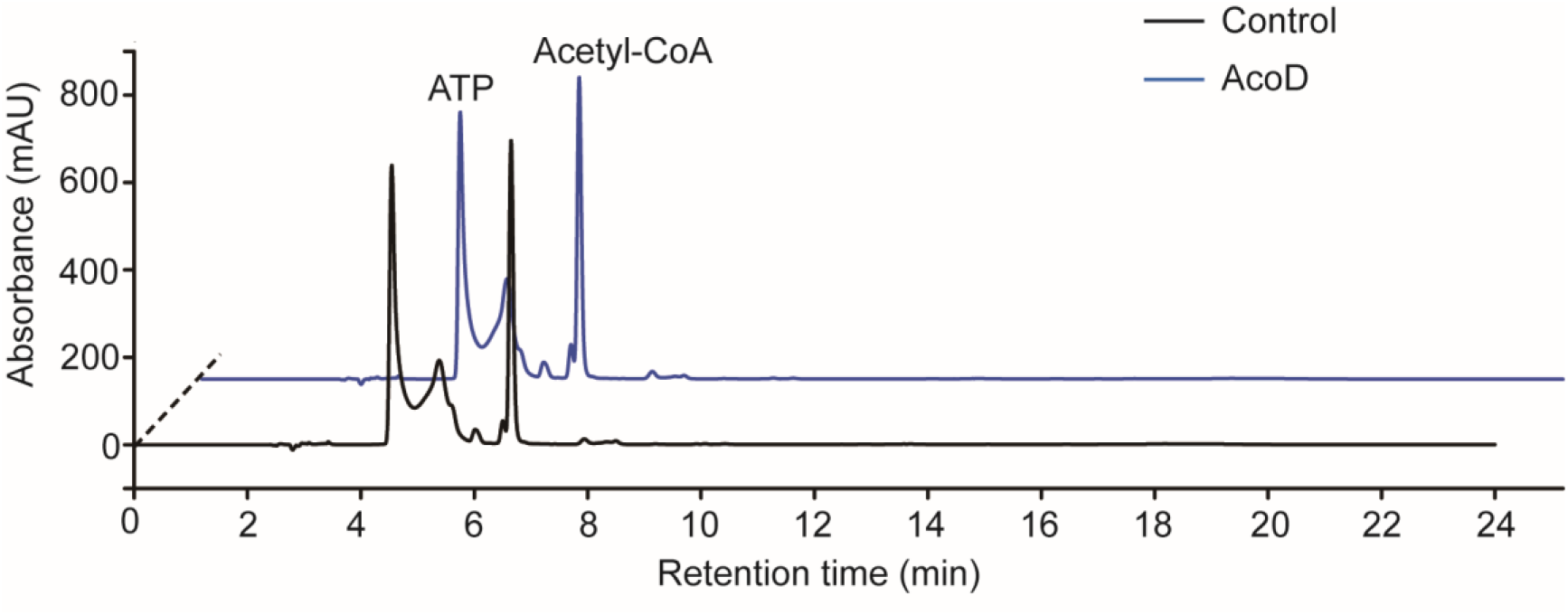
Enzymatic activity analysis of the recombinant AcoD using acetyl-CoA as a CoA donor. ATP and acetyl-CoA were analyzed by HPLC. The result showed that AcoD failed to catalyze the degradation of DMSP when acetyl-CoA was used as a CoA donor.

**Supplementary Table 1.**
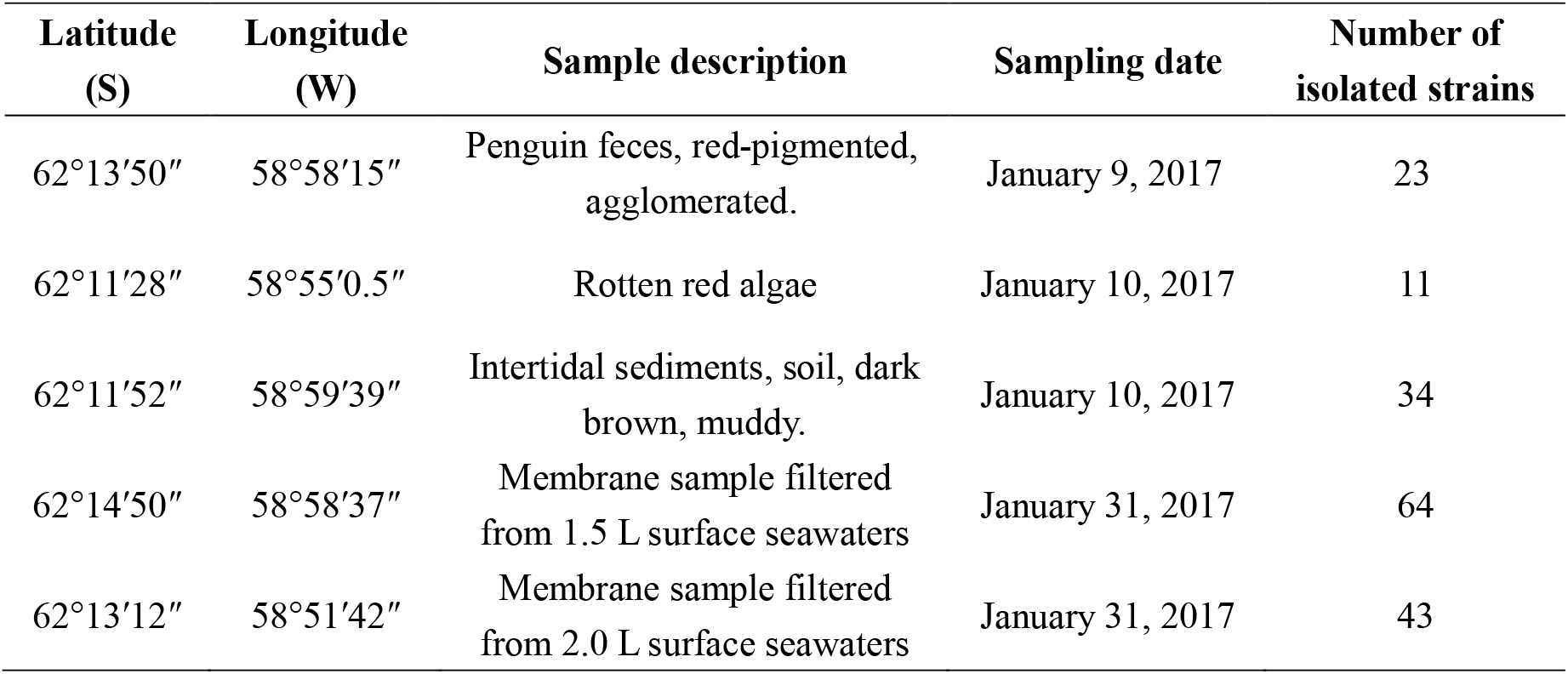
Information of the Antarctic samples used in this study.

| Latitude<br>(S) | Longitude<br>(W) | Sample description | Sampling date | Number of<br>isolated strains |
| --- | --- | --- | --- | --- |
| 62°13'50" | 58°58'15" | Penguin feces, red-pigmented,<br>agglomerated. | January 9, 2017 | 23 |
| 62°11'28" | 58°55'0.5" | Rotten red algae | January 10, 2017 | 11 |
| 62°11'52" | 58°59'39" | Intertidal sediments, soil, dark<br>brown, muddy. | January 10, 2017 | 34 |
| 62°14'50" | 58°58'37" | Membrane sample filtered<br>from 1.5 L surface seawaters | January 31, 2017 | 64 |
| 62°13'12" | 58°51'42" | Membrane sample filtered<br>from 2.0 L surface seawaters | January 31, 2017 | 43 |

**Supplementary Table 2.** Homology alignment of proteins in *Psychrobacter* sp. D2 with known

| Enzymes involved in DMSP catabolism | Proteins of the highest similarity in *Psychrobacter*. sp. D2 | Similarity |
| --- | --- | --- |
| DddD (*Marinomonas* sp. MWYL1) | orf01588 (CoA transferase) | 19.80% |
| DddD (*Ruegeria pomeroyi* DSS-3) | orf01588 (CoA transferase) | 23.62% |
| DddP (*Roseovarius nubinhibens* ISM) | orf00311 (DUF3732 domain-containing protein) | 4.03% |
| DddP (*Roseobacter denitrificans* OCh 114) | orf02289 (Methionine aminopeptidase) | 6.83% |
| DddP (*Ruegeria pomeroyi* DSS-3) | orf01862 (MATE family efflux transporter) | 4.90% |
| DddQ (*Ruegeria lacuscaerulensis* ITI-1157) | orf00350 (1-deoxy-D-xylulose-5-phosphate synthase) | 8.64% |
| DddQ (*Ruegeria pomeroyi* DSS-3) | orf01055 (Rod shape-determining protein RodA) | 7.53% |
| DddY (*Alcaligenes faecalis*) | orf01725 (Nitrite reductase small subunit NirD) | 3.78% |
| DddW (*Ruegeria pomeroyi* DSS-3) | orf02427 (TRAP transporter small permease) | 9.14% |
| DddL (*Rhodobacter sphaeroides* 2.4.1) | orf02192 (dTDP-glucose 4,6-dehydratase) | 6.94% |
| DddL (*Sulfitobacter* sp. EE-36) | orf01537 (Cupin domain-containing protein) | 7.96% |
| DddK (*Candidatus Pelagibacter ubique* HTCC1062) | orf01209 (LysR family transcriptional regulator) | 16.47% |
| Alma1 (*Emiliana huxleyi*) | orf00316 (Molybdenum cofactor biosynthesis protein) | 5.77% |

**Supplementary Table 3.** Crystallographic data collection and refinement parameters of AcoD.

| Parameters | Se-derivative of AcoD | AcoD/ATP complex |
| --- | --- | --- |
| **Diffraction data** |  |  |
| Space group | $P2_12_12_1$ | C2 |
| Unit cell |  |  |
| a, b, c (Å) | 85.3, 87.5, 327.6 | 143.6, 88.8, 273.3 |
| $\alpha, \beta, \gamma$ (°) | 90.0, 90.0, 90.0 | 90.0, 99.9, 90.0 |
| Resolution range (Å) | 50.0-3.3 (3.36-3.30) * | 50.0-2.25 (2.29-2.25) |
| Redundancy | 12.5 (11.6) | 3.2 (3.3) |
| Completeness (%) | 99.8 (98.4) | 97.3 (95.9) |
| $R_{\text{merge}}$ ** | 0.1 (0.4) | 0.1 (0.4) |
| $I/\sigma I$ | 18.0 (3.9) | 13.1 (2.0) |
| **Refinement statistics** |  |  |
| R-factor |  | 0.23 |
| Free R-factor |  | 0.18 |
| RMSD from ideal geometry |  |  |
| Bond lengths (Å) |  | 0.008 |
| Bond angles (°) |  | 1.1 |
| Ramachandran plot (%) |  |  |
| Favored |  | 93.7 |
| Allowed |  | 5.1 |
| Outliers |  | 1.2 |
| Overall B-factors (Å<sup>2</sup>) |  | 42.4 |

\*Numbers in parentheses refer to data in the highest-resolution shell.
\*\* $R_{\text{merge}} = \frac{\sum_{hkl} \sum_i |I(hkl)_i - \langle I(hkl) \rangle|}{\sum_{hkl} \sum_i I(hkl)_i}$ , where $I$ is the observed intensity, $\langle I(hkl) \rangle$
represents the average intensity, and $I(hkl)_i$ represents the observed intensity of each unique
reflection.

**Supplementary Table 4.**
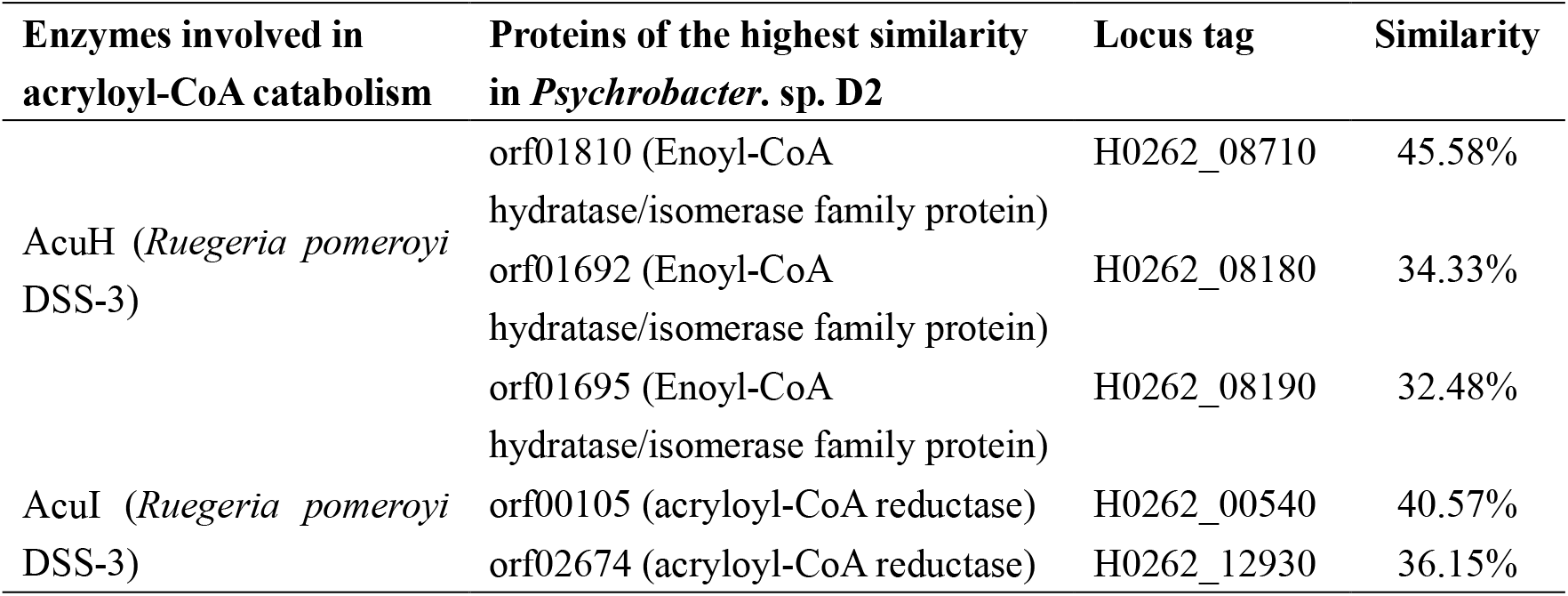
Homology alignment of proteins in *Psychrobacter* sp. D2 with the acryloyl-CoA hydratase AcuH and the acryloyl-CoA reductase AcuI from *Ruegeria pomeroyi* DSS-3.

**Supplementary Table 5.** Strains and plasmids used in this study.

| Strains | Description | Source |
| --- | --- | --- |
| *Psychrobacter* sp. D2 | Wild-type isolate | This study |
| $\Delta acoD$ | the *acoD* gene deletion mutant | This study |
| $\Delta acoD$/pBBR1MCS-*acoD* | $\Delta acoD$ containing pBBR1MCS-*acoD* plasmid | This study |
| $\Delta acoD$/pBBR1MCS | $\Delta acoD$ containing pBBR1MCS plasmid | This study |
| *Escherichia. coli* WM3064 | RP4 (tra) in chromosome, DAP<sup>-</sup>, 37°C | (*Dehio et al., 1997*) |
| *Escherichia. coli* DH5α | Used for gene cloning | Vazyme Biotech company (China) |
| *Escherichia. coli* BL21(DE3) | Used for gene expression | Vazyme Biotech company (China) |
| **Plasmids** |  |  |
| pK18*mobsacB*-Ery | pk18*mobsacB* containing the erythromycin resistant gene from pHT304, Kan<sup>r*</sup>, Ery<sup>r**</sup> | (*Wang et al., 2015*) |
| pK18Ery-*acoD* | pK18*mobsacB*-Ery containing the homologous arms of the *acoD* gene of *Psychrobacter. sp.* D2, Kan<sup>r*</sup>, Ery<sup>r**</sup> | This study |
| pBBR1MCS | Broad-host-range cloning vector, Kan<sup>r*</sup> | (*Kovach et al., 1995*) |
| pBBR1MCS-*acoD* | pBBR1MCS containing the *acoD* gene and its promoter of *Psychrobacter. sp.* D2, Kan<sup>r*</sup> | This study |
| pET-22b | Used for gene expression | Novagen (Germany) |

\* Kan<sup>r</sup>, kanamycin resistance.
\*\* Ery<sup>r</sup>, erythromycin resistance.

**Supplementary Table 6.** Composition of the basal medium (lacking the carbon source).

| Solution* | Components |
| --- | --- |
| 1 | 0.05% (w/v) NH<sub>4</sub>Cl, 3% (w/v) NaCl, 0.3% (w/v) MgCl<sub>2</sub>·6H<sub>2</sub>O, 0.2% (w/v) K<sub>2</sub>SO<sub>4</sub>, 0.02% (w/v) K<sub>2</sub>HPO<sub>4</sub>, 0.001% (w/v) CaCl<sub>2</sub>, 0.0006% (w/v) FeCl<sub>3</sub>·6H<sub>2</sub>O, 0.0005% (w/v) Na<sub>2</sub>MoO<sub>4</sub>·7H<sub>2</sub>O, 0.0004% (w/v) CuCl<sub>2</sub>·2H<sub>2</sub>O, 0.6% (w/v) Tris. [1.5% (w/v) agar for solid medium] |
| 2 | 0.001% (w/v) thiamine·HCl, 0.002% (w/v) nicotinic acid, 0.002% (w/v) pyridoxine·HCl, 0.002% (w/v) riboflavin, 0.0001% (w/v) biotin, 0.0001% (w/v) cyanocobalamin, 0.001% (w/v) *p*-aminobenzoic acid, 0.002% (w/v) calcium pantothenate. |

\* Solution 1 was autoclaved at 121°C for 20 min. Solution 2 was filter-sterilized before it was combined with solution 1.

**Supplementary Table 7.**
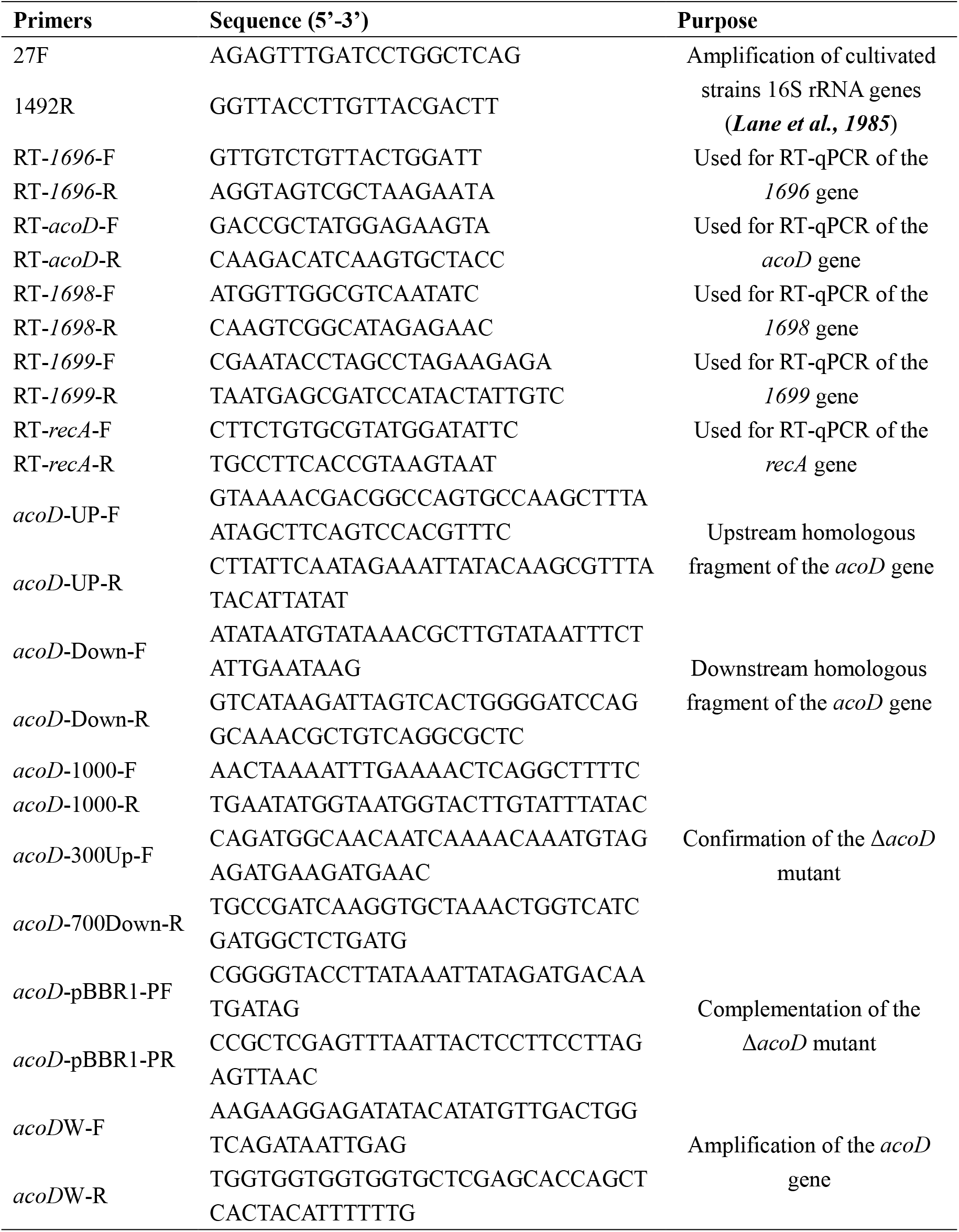
Primers used in this study.

